# Anomalous incisor morphology indicates tissue-specific roles for *Tfap2a* and *Tfap2b* in tooth development

**DOI:** 10.1101/2020.06.18.157776

**Authors:** Emily D. Woodruff, Galaxy C. Gutierrez, Eric Van Otterloo, Trevor Williams, Martin J. Cohn

## Abstract

Mice possess two types of teeth that differ in their cusp patterns; incisors have one cusp and molars have multiple cusps. The patterning of these two types of teeth relies on fine-tuning of the reciprocal molecular signaling between dental epithelial and mesenchymal tissues during embryonic development. Here we show that the incisors are populated only at early time points by the neural crest, whereas the molars continue to receive contributions at later stages, revealing a temporal difference that could alter epithelial-mesenchymal signaling dynamics between these two types of teeth. The AP-2 transcription factors, particularly *Tfap2a* and *Tfap2b*, are essential components of such epithelial-mesenchymal signaling interactions that coordinate craniofacial development in mice and other mammals, but little is known about their roles in the regulation of tooth development and shape. We demonstrate that incisors and molars differ in their temporal and spatial expression of *Tfap2a* and *Tfap2b*; in particular, at the bud stage, *Tfap2a* is expressed in both the epithelium and mesenchyme of the incisors and molars but expression of *Tfap2b* is restricted to the mesenchyme of the molars. Tissue-specific deletions show that loss of the epithelial domain of *Tfap2a* and *Tfap2b* affects the number and spatial arrangement of the incisors, notably resulting in duplicated lower incisors. In contrast, deletion of these two genes in the mesenchymal domain has little effect on tooth development. Collectively these results implicate epithelial expression of *Tfap2a* and *Tfap2b* in dorsal-ventral patterning of the incisors and suggest that these genes contribute to morphological differences between anterior (incisor) and posterior (molar) teeth within the mammalian dentition.

**Highlights:**

1. Late-migrating cranial neural crest cells contribute extensively to the developing molar tooth germs but minimally to the incisors.
2. During tooth development, transcription factors *Tfap2a* and *Tfap2b* are expressed in spatially and temporally dynamic patterns and differ between incisor and molar tooth germs.
3. Epithelial expression of *Tfap2a* and *Tfap2b* is necessary for incisor development, but mesenchymal expression of these genes is not required.

## 1. Introduction

Embryonic development of the tooth crown is divisible into four morphologically distinct stages: initiation, bud, cap, and bell stages (Jernvall and Thesleff, 2012; Tucker and Sharpe, 2004) (Figure 1). Teeth arise from a series of molecular and physical interactions between epithelial and mesenchymal tissues in the oral cavity (Kollar and Baird, 1969; Lumsden, 1988; Mina and Kollar, 1987). The dental epithelium and mesenchyme originate from different embryonic tissues; the dental epithelium is derived from the oral epithelium whereas the dental mesenchyme is derived predominantly from the cranial neural crest cells (CNCCs), with a minor component coming from the head mesoderm (Chai et al., 2000; Douarin and Kalcheim, 1999; Hall, 2009; Lumsden, 1988, 1987; Mina and Kollar, 1987; Rothová et al., 2011). Mutations that affect these tissue interactions (*e.g., Pitx2, Msx1,* and *Pax9*) can cause profound disruptions to the development of the human dentition, including loss, gain, or mis-patterning of teeth (Alappat et al., 2003; Chen et al., 1996; Dressler et al., 2010; Mostowska et al., 2003; Peters et al., 1998; Satokata and Maas, 1994).

**Figure 1.**
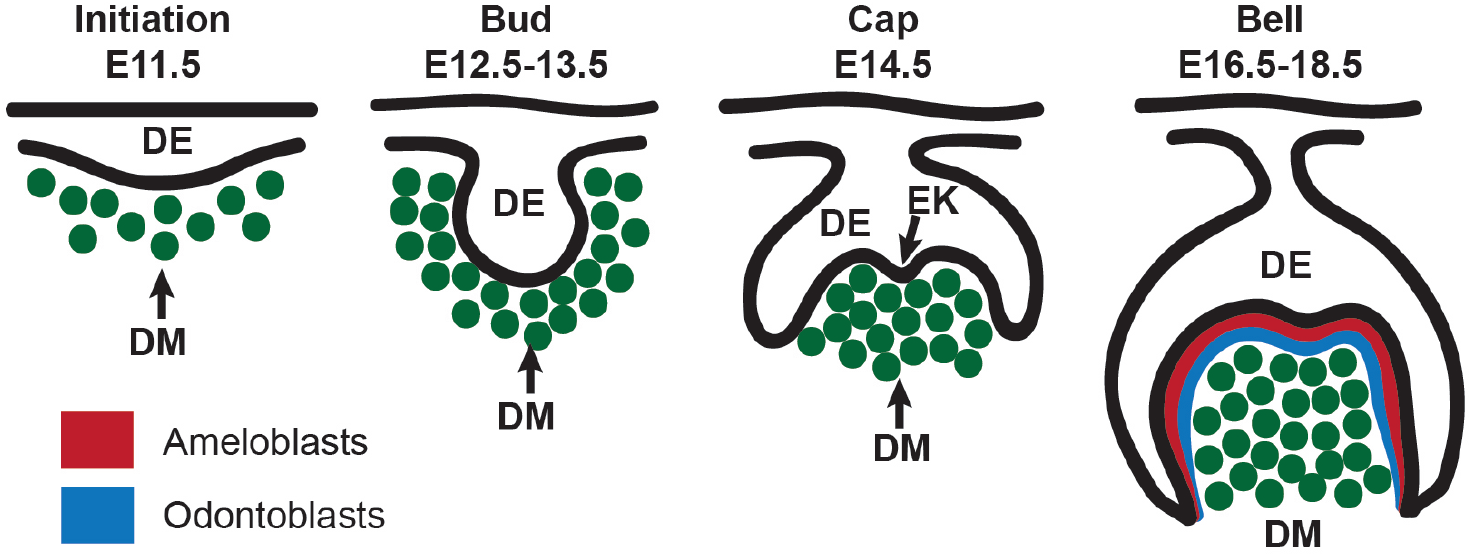
Stages of mouse tooth development illustrating epithelial and mesenchymal tissues in a lower molar tooth germ at the initiation, bud, cap, and bell stages. Once tooth development is initiated, the dental epithelium grows into the adjacent dental mesenchyme which condenses around the epithelial bud. During the cap stage the epithelium then grows around the condensed mesenchyme and the primary enamel knot forms which initiates the patterning of the tooth cusp(s). Cytodifferentiation takes place during the bell stage when the ameloblasts and odontoblasts form which will deposit enamel and dentine, respectively. DE: dental epithelium, DM: dental mesenchyme, EK: enamel knot. Drawing after Tucker and Sharpe, 2004.

In mice, regionalization of the dentition into presumptive incisor and molar domains occurs between embryonic day (E) 9 and E10, when CNCCs, which comprise the majority of the branchial arch mesenchyme, reach the end of their migration into the first branchial arch (Lumsden, 1988; Chai et al., 2000). During the bud stage (E12.5-E13.5), the dental epithelium proliferates into the dental mesenchyme, which condenses around the epithelial bud (Figure 1). In the cap stage (E14.5), the dental epithelium continues to proliferate around the condensed dental mesenchyme cells, and the primary enamel knot appears at the center of the invaginating epithelial bud during the transition from bud to cap stages (E13.5-E14.5) (Cho et al., 2007; Jernvall et al., 1994; Vaahtokari et al., 1996). The enamel knot, a transient localized cluster of non-proliferating epithelial cells, is an important signaling center that controls tooth shape, specifically the pattern of cusps on the tooth (Jernvall et al., 1994; Vaahtokari et al., 1996). In teeth with multiple cusps, the primary enamel knot is thought to direct the formation of other subsequent enamel knots that are associated with individual tooth cusps (Jernvall et al., 1994; Jernvall and Thesleff, 2012; Luukko et al., 2003; Thesleff et al., 2001). The bell stage (E16.5-18.5) is the final stage in embryonic dental development in which the occlusal cusp pattern takes its final shape, molecularly determined by the enamel knots and the folding of the inner enamel epithelium. In the inner enamel epithelium, differentiation ensues and the resulting ameloblasts secrete enamel matrix, while dental mesenchymal cells differentiate into odontoblasts which form the dentine and inner pulp (Nanci, 2013).

In addition to initiating odontogenesis and inducing the cell types that produce dental tissues, epithelial-mesenchymal tissue interactions are essential for the formation of morphologically distinct tooth types within the dentition (Kollar and Baird, 1969; Lumsden, 1988; Mina and Kollar, 1987). Most mammals, including humans, possess multiple differently-shaped teeth which are associated with specific dietary specializations in some species. Mice, for example, have two different types of teeth in their dentition: two anterior upper and lower incisors, which each have a single cusp, and six posterior upper and lower molars which are multi-cusped. The genetic basis for tooth development has been well-studied in mice, with considerable emphasis on the formation of cusps on the molar crown (Ahn et al., 2010; Harjunmaa et al., 2012; Jernvall et al., 1994; Pispa et al., 1999; Thesleff et al., 2001) and the ever-growing properties of murine incisors (Harada et al., 2002; Klein et al., 2008; Tummers and Thesleff, 2003). Less attention has been paid, however, to incisor crown formation, and relatively few studies have explicitly compared gene expression patterns between developing incisors and molars (Hu et al., 2013; Huang et al., 2014; Laugel-Haushalter et al., 2013; Tucker et al., 1998b).

Here we used an inducible lineage tracing approach in mice and uncovered striking differences in the timing of neural crest cell contribution between incisors and molars. To investigate these differences further, we compared the molecular identities of incisors and molars in the context of two AP-2 paralogs, *Tfap2a* and *Tfap2b*. Both genes are members of the activator protein-2 (AP-2) family of transcription factors (Williamson et al., 1996), of which five AP-2 proteins are known in mammals, AP-2*α*-AP-2ε (Eckert et al., 2005). AP-2 transcription factors are expressed early in development in the neural crest and they are known to play an essential role in craniofacial development in numerous vertebrate species, including mice, zebrafish, and chickens (Brewer et al., 2004; Brewer and Williams, 2004; de Croze et al., 2011; Hoffman et al., 2007; Knight et al., 2005; Li and Cornell, 2007; Mitchell et al., 1991; Nottoli et al., 1998; Van Otterloo et al., 2018; Zhang et al., 1996); however, little is known about their roles in tooth development. Recent transcriptional profiling studies of developing teeth in mice (Laugel-Haushalter et al., 2013), humans (Huang et al., 2014), and minipigs (Wang et al., 2014) have identified expression of many genes, including *Tfap2a* (Wang et al., 2014) and *Tfap2b* (Laugel-Haushalter et al. 2013; Wang et al., 2014). Expression of *Tfap2b* was also reported in the dental mesenchyme of mouse molars at bud-bell stages (Tanasubsinn et al., 2017; Uchibe et al., 2012) contradicting results from an earlier study in which *Tfap2a* was detected in tooth germs but *Tfap2b* was reported as absent (Moser et al., 1997). These earlier investigations of AP-2 expression in teeth were limited to molars and results were not reported for incisors, with the exception of one study that showed a lack of *Tfap2b* expression in murine upper incisors (Tanasubsinn et al., 2017). Finally, several human genetic studies have identified dental anomalies in patients with *Tfap2a* and *Tfap2b* mutations, which cause the human syndromic disorders branchio-oculo-facial syndrome and Char syndrome, respectively (Milunsky et al., 2008; Satoda et al., 2000; Tanasubsinn et al., 2017). The relevance of *Tfap2a* and *Tfap2b* to human disease underscores the need for a detailed characterization of the expression domains and tissue-specific functions of AP-2 genes during dental development.

To this end, we compared spatiotemporal differences in the expression of both *Tfap2a* and *Tfap2b* between incisors and molars throughout tooth development and used mouse conditional genetics to determine the tissue-specific roles of these genes in dental epithelium and mesenchyme of each tooth class. Though *Tfap2a* and *Tfap2b* are expressed in epithelial and mesenchymal tissues during tooth development, we find that epithelial-specific loss of *Tfap2a* and *Tfap2b* resulted in a loss or reduction of upper incisors along with a duplication of lower incisors, but deletion of these genes in the neural crest-derived mesenchyme did not perturb dental development. Despite major impacts on incisor development following epithelial loss of *Tfap2a* and *Tfap2b,* molar development was essentially unaffected. Collectively, our results identify a novel role for *Tfap2* family members in dental development.

## 2. Materials and Methods

### 2.1 Mice

All animal procedures were conducted under strict accordance of all applicable guidelines and regulations, following the ‘Guide for the Care and Use of Laboratory Animals of the National Institutes of Health’. In addition, all animal experiments conducted were approved by the Institutional Animal Care and Use Committees of the University of Florida or the University of Colorado – Denver, depending on the mouse line used (further outlined below).

ICR (CD-1) “wild-type” laboratory mice (Envigo) were used for histology and *in situ* hybridization experiments. Two transgenic strains (Jackson Laboratory) were used to perform lineage tracing studies on neural crest cells, *CBA;B6-Tg(Sox10-icre/ER*^*T2*^)^*388Wdr/J*^ (JAX stock number 027651, abbreviated *Sox10-iCre/ER^T2^*) (McKenzie et al., 2014), and *Gt(ROSA)26Sor*^*tm4(ACTB-tdTomato,-EGFP)Luo*^ (JAX stock number 007576, abbreviated *R26R*^*mTmG*^) (Muzumdar et al., 2007). CD-1, *Sox10-iCre/ER*^*T2*^, and *R26R*^*mTmG*^ mice were housed in the University of Florida Animal Care Services barrier facility. All *Tfap2a* and *Tfap2b* mutant lines were housed at the University of Colorado-Denver. Mice had access to food and water *ad libitum*. For all timed matings, the day on which a copulatory plug was detected in the female, the embryos were denoted as E0.5.

At the appropriate developmental stage, embryos were collected by first euthanizing the pregnant dam, dissecting out the uterine horn, removing the embryos from the uterine muscle and extraembryonic tissue in ice-cold phosphate buffered saline (PBS) or PBS treated with diethyl-pyrocarbonate (DEPC-PBS). Embryos were staged according to embryonic days as previously described (Kaufman, 1992; Martin, 2002) and processed in a manner contingent on their downstream application (described below). A small portion of the extraembryonic yolk-sac or tail snippet of the dissected embryo was saved for DNA extraction and genotyping. All adult mice and embryos in this study were genotyped using polymerase chain reaction (PCR) (see Supplementary Table 1 for primer sequences).

### 2.2 Lineage tracing of *Sox10*-expressing neural crest cells

To follow cranial neural crest cells (CNCCs) and their derivatives from their original location in the neural crest to their final destinations in the face and jaw, including the teeth, we crossed male mice heterozygous for a tamoxifen-inducible *Cre* allele driven by the *Sox10* promoter (*Sox10-iCre/ER^T2^*) (McKenzie et al., 2014) with females heterozygous or homozygous for double-fluorescent *Cre* reporter alleles (*R26R*^*mTmG/wt*^ or *R26R*^*mTmG/mTmG*^) (Muzumdar et al., 2007) (see Supplementary Methods 5.1 for details). In embryos from this cross, *Sox10-iCre*-positive CNCCs and their derivatives were EGFP-positive. Migrating CNCCs express *Sox10* but, *Sox10* is not expressed in the head mesoderm or facial epithelium during this time (Anderson et al., 2006; Britsch et al., 2001; Jacques-Fricke et al., 2012; McKenzie et al., 2014; Ota et al., 2004; Soo et al., 2002), making it a suitable genetic marker for labeling CNCCs in the teeth.

In the first lineage tracing experiment, pregnant female mice were administered tamoxifen (see Supplementary Methods 5.1) to induce *Cre*-mediated recombination on one day when the embryos were at stage E6.5, E7.5, or E8.5. In the second experiment, tamoxifen was administered on three consecutive days from E6.5-8.5. Results are based on a minimum of 3 embryos for each time point in each experiment. Membrane fluorescence for EGFP and tdTomato was visualized in frontal cryosections using a Zeiss LSM 710 confocal microscope.

### 2.3 Conditional Deletion of *Tfap2a* and *Tfap2b*

To generate *Tfap2* mutant embryos, we used conditional floxed or null alleles of *Tfap2a* and *Tfap2b* and two strains in which *Cre* recombinase was expressed in either the epithelium, *Crect* (Schock et al., 2017), or the neural crest, *Wnt1-Cre* (Danielian et al., 1998). Males were heterozygous for either the epithelial or neural crest *Cre* allele and the *Tfap2a* and *Tfap2b* conditional alleles (*i.e., Tfap2a^flox/wt^;Tfap2b^flox/wt^;Wnt1-Cre* or *Tfap2a^flox/wt^;Tfap2b^flox/wt^;Crect)* and females were homozygous for the conditional alleles (*i.e., Tfap2a^flox/flox^*;*Tfap2b^flox/flox^*). Both conditional alleles have been previously described, including, *Tfap2a*^*tm2Will/J*^ (the *Tfap2a* floxed conditional allele) (Brewer et al., 2004) and *Tfap2b*^*tm2Will*^ (the *Tfap2b* floxed conditional allele) (Martino et al., 2016; Seberg et al., 2017; Van Otterloo et al., 2018). In the second cross, males were heterozygous for conditional null alleles of *Tfap2a* (Zhang et al., 1996), and *Tfap2b* (Martino et al., 2016; Seberg et al., 2017; Van Otterloo et al., 2018), (*i.e., Tfap2a^null/wt^;Tfap2b^null/wt^;Wnt1-Cre* or *Tfap2a*^*null/wt*^;*Tfap2b*^*null/wt*^;*Crect*) and females were homozygous for the conditional alleles. In double mutant embryos from the first cross (*i.e., Tfap2a^flox/flox^;Tfap2b^flox/flox^;Cre+*), *Tfap2a* and *Tfap2b* were deleted in the *Cre*-positive tissue (epithelium or CNC-derived mesenchyme) (Supplementary figure 1 A, B). In the second cross, double mutant embryos (*i.e., Tfap2a^null/flox^;Tfap2b^null/flox^;Cre+*) lacked both alleles of *Tfap2a* and *Tfap2b* in the *Cre*-positive tissue and were heterozygous for *Tfap2a* and *Tfap2b* in the rest of the embryo (Supplementary figure 1 C, D) (see Supplementary Methods 5.2 for additional details on tissue-specific deletions). For embryos examined from both *Crect* and *Wnt1-Cre* crosses, genotypes, phenotypes, and sample sizes are provided in Supplementary table 2.

### 2.4 Tissue preparation, cryosectioning, and histology

Tissue was prepared for cryosectioning as follows: mouse heads were fixed overnight in 4% PFA at 4°C, equilibrated in 15% – 30% sucrose in PBS on ice for ~3 hours or overnight depending on the stage, and placed in a solution with equal amounts of OCT and sucrose (30% sucrose) at 4°C overnight. Heads were embedded in OCT, frozen on dry ice, and stored at −80°C. 10μm sections in the frontal (coronal) plane were cut using a Leica cryostat and mounted on Superfrost Plus Gold slides (Thermo Fisher Scientific) (for *in situ* hybridization only) or on Superfrost Plus slides (Thermo Fisher Scientific) and stored at −80°C. Histological staining was performed on cryosections using 10% neutral buffered formalin to post-fix tissue, Mayer’s hematoxylin (Electron Microscopy Sciences), Eosin-Y alcoholic (Fisher Scientific), and Scott’s solution (10g/L MgSO_4_ + 2g/L NaHCO_3_ + tap water). Sections were dehydrated with ethanol, then Xylene, mounted with Permount (Fisher Scientific), and covered with glass coverslips (Thermo Fisher Scientific).

### 2.5 Design and cloning of RNA probes for *in situ* hybridization

Mouse (*Mus musculus*) messenger RNA (mRNA) sequences for genes of interest (*Tfap2a, Tfap2b, Yeats4, Kctd1, Ets1*) were obtained from NCBI GenBank (http://www.ncbi.nlm.nih.gov/genbank/). Oligonucleotide primers (Supplementary table 3) were designed in Geneious (v6.1.8 or 10.0.9, Biomatters, Ltd) and target sequences were PCR-amplified from cDNA or genomic DNA from CD-1 mice, ligated into vectors, and cloned (see Supplementary Methods 5.2 for details). “Sense” (negative control) and “antisense” (experimental) digoxigenin (DIG)-labeled RNA probes were synthesized from the target DNA, purified, and quantified (see Supplementary Methods 5.3).

### 2.6 RNA *in situ* hybridization on cryosections

*In situ* hybridization (ISH) was performed as described previously (Acloque et al., 2008) with some modifications (see Supplementary Methods 5.4 for a complete description). Expression patterns reported here for each gene of interest were detected in a minimum of 3 embryos per stage. The expression patterns of these genes had been previously documented in some region in the head (often in the brain or the eye) and these tissues/regions were used as positive controls (Supplementary figure 2 A-G). Negative controls were also conducted for each gene of interest (Supplementary figure 2 H-L).

### 2.7 Micro-CT scanning and 3-D reconstruction of Tfap2 mutant and control embryos

Mouse embryos were prepared for micro-CT (μCT) using Lugol’s iodine solution for contrast-enhancement (see Supplementary Methods 5.5). Embryos were scanned in a GE V|TOME|X M 240 Nano CT scanner (General Electric) at the University of Florida Nanoscale Research Facility. Tiff stacks were generated using Phoenix Datos2 software (General Electric) and VG Studio Max (Volume Graphics) was used for 3-D reconstructions.

### 2.8 Quantification and analysis of molar occlusal dimensions

The first upper and lower molar teeth (M^1^/^1^) from both left and right sides were measured in VG Studio Max (5 replicate measurements/tooth). Length:width ratios were compared to previously published measurements collected from wild populations of mice, *Mus musculus musculus* (Csanady and Mosansky, 2018; Wallace, 1968). The use of ratios in these comparisons eliminates the effect of differences in overall tooth size. The molar ratios of the control embryos were within the range of those previously calculated for wild mice (Csanady and Mosansky, 2018), and, lacking replicate 3-D data sets, we reasoned that in a larger sample of our laboratory mouse strains, the variance would be similar to that of wild *M. musculus*. Based on this assumption, we used the largest standard deviation from the wild-type data set (+/− 0.08, N=101) (Csanady and Mosansky, 2018) to conservatively estimate hypothetical distributions of length and width measurements for both control and mutant mice from *Crect* and *Wnt1-Cre* crosses (Supplementary table 4). Distributions composed of 100 molar length and width measurements were generated in R (v3.6.0) using the *runif* function, which pseudo-randomly generates values between specific minimum and maximum values. Molar ratios (length:width) were calculated from these distributions and the data were tested for normality using the Shapiro-Wilks test (*shapiro.test* in R). On account of the data being non-normally distributed, the Wilcox-signed rank test (*wilcox-test* in R) was employed to compare simulated molar ratios between the mutant and control embryos.

## 3. Results

### 3.1 Cranial neural crest cells complete their migration to the incisor region prior to the molar region

It is well understood that CNCCs populate the embryonic dental mesenchyme just prior to the initiation of tooth development and that they are important for epithelial-mesenchymal signaling (Chai et al., 2000; Douarin and Kalcheim, 1999; Hall, 2009; Imai et al., 1996; Lumsden, 1988, 1987; Mina and Kollar, 1987; Rothová et al., 2011) but the timing of CNCCs’ arrival into the presumptive incisor (anterior) and molar (posterior) regions is unknown. To address this, we performed lineage tracing of *Sox10-*expressing CNCCs into the developing incisor and molar teeth. Based on previous work (Imai et al., 1996) we hypothesized that the anterior odontogenic (future incisor) region would be colonized by CNCCs before the more posterior (molar) odontogenic region. *Sox10*-positive cells were labeled using a tamoxifen-inducible transgenic mouse line, *Sox10-iCre/ER*^*T2*^ (McKenzie et al., 2014), crossed with an *R26R*^*mTmG*^ fluorescent reporter (Muzumdar et al., 2007). Migration of CNCCs begins at approximately E7.5, therefore, we administered tamoxifen at E6.5, E7.5, or E8.5 to induce *Cre*-recombination in the embryos to label early and late migrating neural crest cells.

In *Cre*-positive embryos from mice that were given tamoxifen at E6.5 or E7.5, EGFP-positive neural crest cells migrated into the mesenchyme of both incisors and molars and make up the majority of the dental mesenchyme at the late bud stage (E13.5) (Figure 2 A, B, D, E, G, H). Contrary to our expectations, in embryos exposed to tamoxifen at E6.5, EGFP-positive CNCCs were broadly distributed in both incisors and molars (Figure 2 A, D, G), implying that early-migrating crest cells contribute concurrently to both anterior and posterior tooth germs. In contrast to these early-migrating CNCCs, the contribution of late-migrating cells (labeled with tamoxifen at E8.5) to the incisors is minimal (Figure 2 C, F), but many late-migrating neural crest cells were present in the molar mesenchyme (Figure 2 I). These results show that late-migrating CNCCs contribute to the molar mesenchyme, but relatively few of these cells migrate further anteriorly into the incisor mesenchyme. In embryos from female mice given three consecutive doses of tamoxifen (E6.5-E8.5), which labels crest cells throughout the majority of their migratory period, results were similar to those of the single day labeling at E6.5 and E7.5 (Supplementary figure 3), and no differences were observed between tooth types in the three-day labeling experiment. Collectively, the results of the lineage tracing analyses suggest that early-migrating (*i.e.,* E6.5, E7.5) CNCCs contribute to both incisor and molar mesenchyme but late-migrating (*i.e.,* E8.5) CNCCs continue to populate the molar mesenchyme.

**Figure 2.**
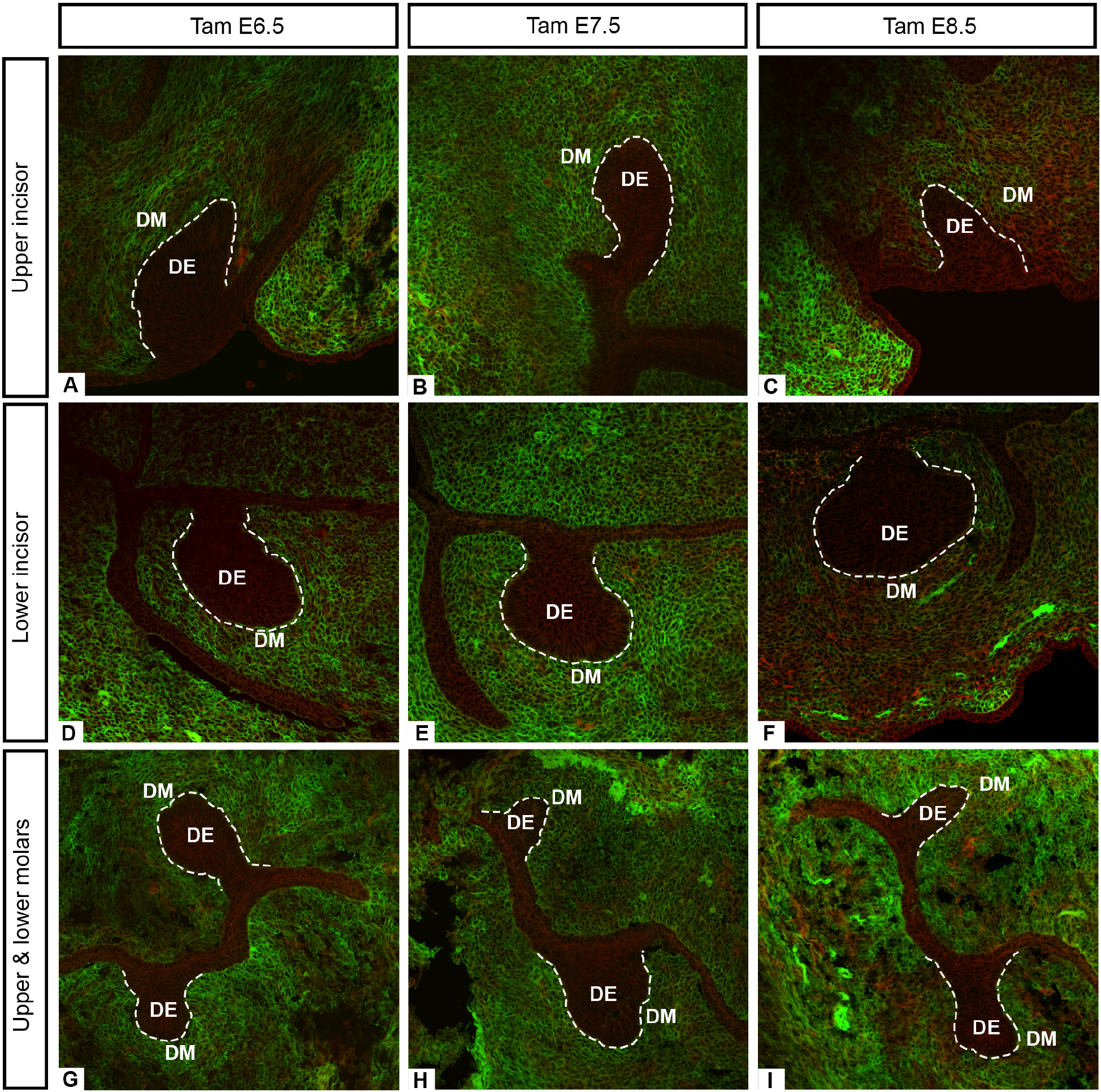
CNCC-derived mesenchyme cells (EGFP-positive) labeled via tamoxifen administration at E6.5 (A, D, G), E7.5 (B, E, H), or E8.5 (C, F, I) in *Sox10-iCre/ER^T2^;R26R^mTmG/+^* or *Sox10-iCre/ER^T2^;R26R^mTmG/mTmG^* embryos. Examination of bud stage tooth germs (E13.5) revealed that fewer CNCC-derived mesenchyme cells were present in the incisor mesenchyme at E8.5 (C, F) compared to the molar mesenchyme (I) seen here in representative frontal cryosections. The dental epithelium is outlined in white. Images taken at 20X. DE: dental epithelium, DM: dental mesenchyme.

### 3.2 Incisors and molars differ in temporal and spatial expression of *Tfap2a* and *Tfap2b*

To further investigate neural crest contributions to incisor and molar development, we chose to study the expression of two AP-2 genes, *Tfap2a* and *Tfap2b* because this transcription factor family is part of the regulatory network driving neural crest development in many species (Mitchell et al., 1991; Moser et al., 1997; Sauka-Spengler and Bronner-Fraser, 2008; Simões-Costa and Bronner, 2015). To determine the spatial and temporal expression of *Tfap2a* and *Tfap2b* we compared mRNA localization in mouse incisors and molars at E12.5-E13.5 (bud stage), E14.5 (cap stage), and E16.5 (early bell stage) using *in situ* hybridization.

*Tfap2a* expression was detected throughout the bud and cap stages in the dental epithelium and mesenchyme of both the incisor and the molar buds (Figure 3 A-C, G-I, M-O). In contrast, *Tfap2b* was not detected in the incisors (epithelium or mesenchyme) until the late bud stage (E13.5) when minimal expression was observed in the incisor epithelium (Figure 3 D-E, J-K), and in some embryos expression was not detected, suggesting that at E13.5 *Tfap2b* transcripts were just beginning to accumulate in the incisor buds. In the molar buds, however, *Tfap2b* was expressed in the mesenchyme during this time (E12.5-E13.5) (Figure 3 F, L), and this expression persisted in the cap stage molars (Figure 3 R), consistent with previous reports (Tanasubsinn et al., 2017; Uchibe et al., 2012). At the cap stage *Tfap2b* was also faintly detected in the incisors (Figure 3 P, Q), the oral and facial epithelia, and the mesenchyme surrounding the nasal cavity (Figure 3 P, J; Supplementary figure 2 E, G).

**Figure 3.**
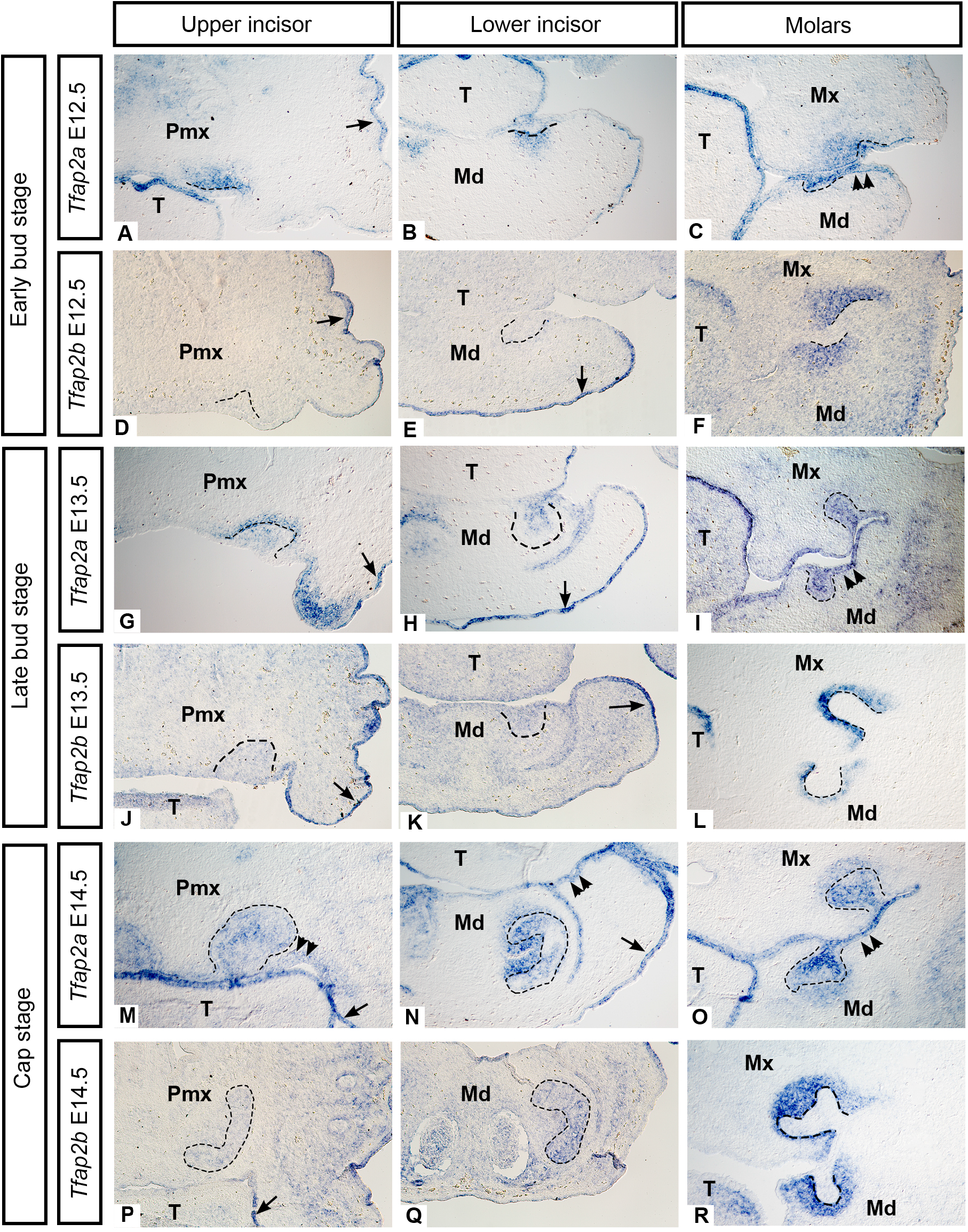
Bud stage (E12.5 and 13.5) and cap stage (E14.5) mRNA expression of *Tfap2a* and *Tfap2b.* mRNA transcripts detected by *in-situ* hybridization on frontal cryosections through the upper incisor (left column), lower incisor (middle column), and molars (right column) are shown and the dental epithelium is outlined. There is minimal expression of *Tfap2b* in the bud stage upper and lower incisors (D-E, J-K) compared to the molar buds (F, L). Both *Tfap2a* and *Tfap2b* were detected in the surface epithelium (arrows) but only *Tfap2a* was present in the oral epithelium (double arrowheads). Note that only the right or left side of each frontal section is shown. Images taken of the embryos’ right side (B, C, H, I, J, K, P, Q) have been mirrored to match images taken of the left side. All images taken at 10X magnification. Pmx: premaxilla, Mx: maxilla, Md: mandible, T: tongue.

In contrast to earlier stages, few *Tfap2a* transcripts were detected in the lower incisor epithelium at the early bell stage (E16.5) (Figure 4 E) while in the upper incisor *Tfap2a* was expressed in the epithelium and the mesenchyme (Figure 4 A). In bell stage molars, however, *Tfap2a* expression was restricted to the inner enamel epithelium directly adjacent to the dental mesenchyme (Figure 4 I, M). In early bell stage incisors, *Tfap2b* transcripts were detected prominently within epithelial-derived ameloblasts and faintly in the mesenchyme (Figure 4 B, F); however, in the molars, *Tfap2b* expression became limited to mesenchymal cells closer to the outer regions of the tooth germ (Figure 4 J, N). At all stages examined, *Tfap2a* was also prominently expressed in the oral epithelium and/or surface epithelium (Figure 3 A-C, G-I, M-O; Figure 4 A, E), in agreement with previous studies (Zhang and Williams, 2003; Zhao et al., 2011).

**Figure 4.**
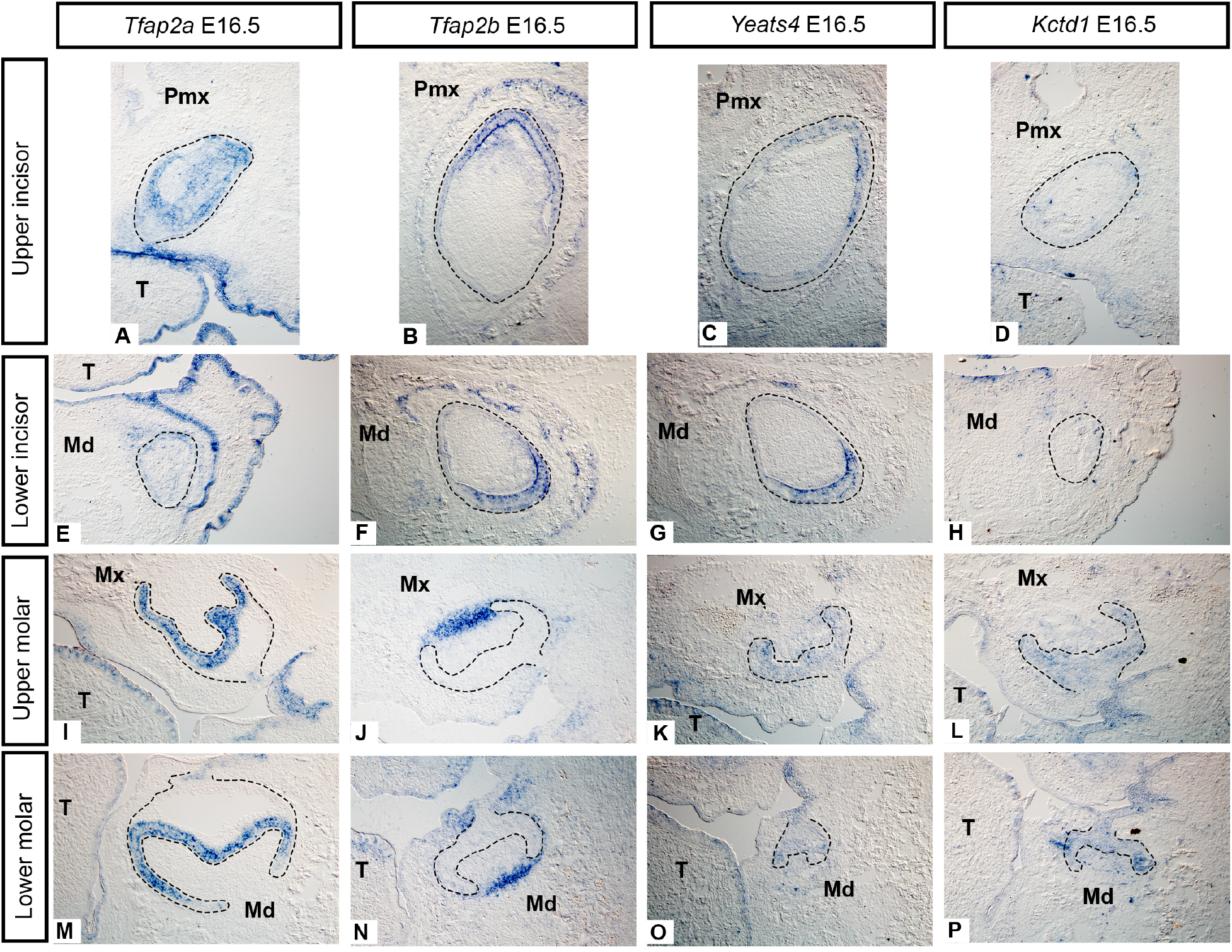
Early bell stage (E16.5) mRNA expression of *Tfap2a, Tfap2b, Yeats4,* and *Kctd1.* mRNA transcripts detected by *in-situ* hybridization on cryosections in the frontal plane are shown in the upper incisor (A-D), lower incisor (E-H), upper molar (I-L), and lower molar (M-P). The dental epithelium is outlined. Note in particular the highly restricted epithelial expression domain of *Tfap2a* in the molars (I, M) and the overlapping expression domains of *Tfap2b* and *Yeats4* in similar regions within the upper (B, C) and lower incisors (F, G). Note that only the right or left side of each frontal section is shown. Images taken of the embryos’ right side (C, F, G, I, M, N, J, K, O) have been mirrored to match images taken of the left side. All images taken at 10X magnification. Pmx: premaxilla, Mx: maxilla, Md: mandible, T: tongue.

We next asked whether *Tfap2a* and *Tfap2b* expression in molars and incisors is associated with *Yeats4*, which encodes an AP-2 activator protein (Ding et al., 2006) and *Kctd1* which encodes an AP-2 inhibitor (Ding et al., 2009). *Yeats4* transcripts were detected in the epithelium of both the incisor and molar buds at E13.5 (Supplementary figure 4 A-C). *Kctd1* was expressed robustly in bud-stage upper incisor epithelium and to a lesser extent in the mesenchyme but minimal expression was detected in the lower incisors (Supplementary figure 4 D, E). In the molars at E13.5, *Kctd1* transcripts are present in the epithelium (Supplementary figure 4 F). By the cap stage, transcripts of both *Yeats4* and *Kctd1* were detected in epithelium and mesenchyme within the incisors and molars (Supplementary figure 4 G-I, J-L) similar to *Tfap2a* and, to a lesser extent, *Tfap2b* (mesenchyme only).

At the early bell stage, *Yeats4* expression was restricted to the incisors where it was localized to epithelium-derived ameloblasts (Figure 4 C, G), similar to that of *Tfap2b*, but unlike *Tfap2b* it was also detected in the epithelium in the molars (Figure 4 K, O). In contrast, *Kctd1* expression was visible in only a few cells in early bell stage incisors and some embryos showed no staining at this stage (Figure 4 D, H). In the molars, however, epithelial expression of *Kctd1* persisted but mesenchymal expression was no longer detectable (Figure 4 L, P).

We then compared these patterns with *Ets1* which, like *Tfap2a* and *Tfap2b* is also expressed in migrating CNCCs where it has been shown to act downstream of *Tfap2a* in the chick (Barembaum and Bronner, 2013). *Ets1* expression was detected in the mesenchyme at cap and early bell stages in the incisors (Supplementary figure 5A, B, E, F) and in the molars (Supplementary figure 5 C, G, H) but its expression was punctate, particularly in the molars. At the early bell stage *Ets1* transcripts were also observed in the dental epithelium. Comparison of *Ets1* expression with comparable histological sections revealed similarities between *Ets-1*-expressing cells and erythrocytes with respect to cell shape and distribution within the tooth germ (Supplementary figure 5 D).

In summary, *Tfap2a, Kctd1,* and *Yeats4* exhibit similar expression patterns at bud-cap stages (E13.5 and E14.5), particularly in the dental epithelium of the incisors and molars, however, expression of *Tfap2b* is similar to the others only at E14.5 in the incisor epithelium and the molar mesenchyme. By the early bell stage (E16.5) expression patterns of these genes differ considerably from one another with the exception of *Tfap2b* and *Yeats4* which are expressed in similar patterns in the ameloblast layer in the incisors. We did not detect similar expression patterns for *Ets1* and *Tfap2a*/*Tfap2b* in incisors or molars. In all ISH assays negative (sense) controls for each probe produced no signal (Supplementary figure 2 H-L).

### 3.3 Epithelial deletion of *Tfap2a* and *Tfap2b* leads to misshapen teeth and extra incisors

The dynamic expression patterns of both *Tfap2a* and *Tfap2b* in the dental epithelium and mesenchyme suggested that these factors could play tissue-specific roles in tooth development. To test whether epithelial-specific expression of *Tfap2a* and *Tfap2b* was required for the development of properly shaped teeth, we used an epithelial-specific *Cre* recombinase allele, *Crect* (Schock et al., 2017). Mutant embryos were homozygous for *Tfap2a* and *Tfap2b* conditional alleles and heterozygous for the *Crect* transgene (*Tfap2a^flox/flox^;Tfap2b^flox/flox^;Crect*), resulting in deletion of *Tfap2a* and *Tfap2b* exclusively from the ectoderm, including the presumptive dental epithelium (Supplementary figure 1 A).

In E18.5 control embryos, hemi-mandibles possess a single upper and single lower incisor (I^1^/_1_) and first and second upper and lower molars (M^1^/_1_, M^2^/_2_) (Figure 5 A, C, F, I, K; Supplementary figure 6 F). By contrast, the most striking difference in E18.5 embryos lacking *Tfap2a* and *Tfap2b* in the epithelium were changes in the number and/or morphology of the lower incisors (Supplementary table 2). In some instances, the mutants displayed an additional lower incisor ventral to I_1_ (N=2/4 embryos) (Figure 5 B). Note that this phenotype was consistently observed in both the left and right hemi-mandibles of the mutants. Morphologically these additional teeth looked similar to I_1_ and histological analysis revealed that a complete repertoire of differentiated cell types were present in this supernumerary incisor, including enamel-forming ameloblasts and dentin-forming odontoblasts (Figure 5 B, B’). μCT scanning and subsequent 3-D reconstruction and histological analysis showed that the mutants that lacked duplicated incisors had an aberrantly shaped lower incisor (I_1_) that exhibited ventral curvature (N=2/4 embryos) (Figure 5 M, O; Supplementary figure 7 C-D); this phenotype was also bilaterally symmetrical and was observed in both right and left hemi-mandibles.

**Figure 5.**
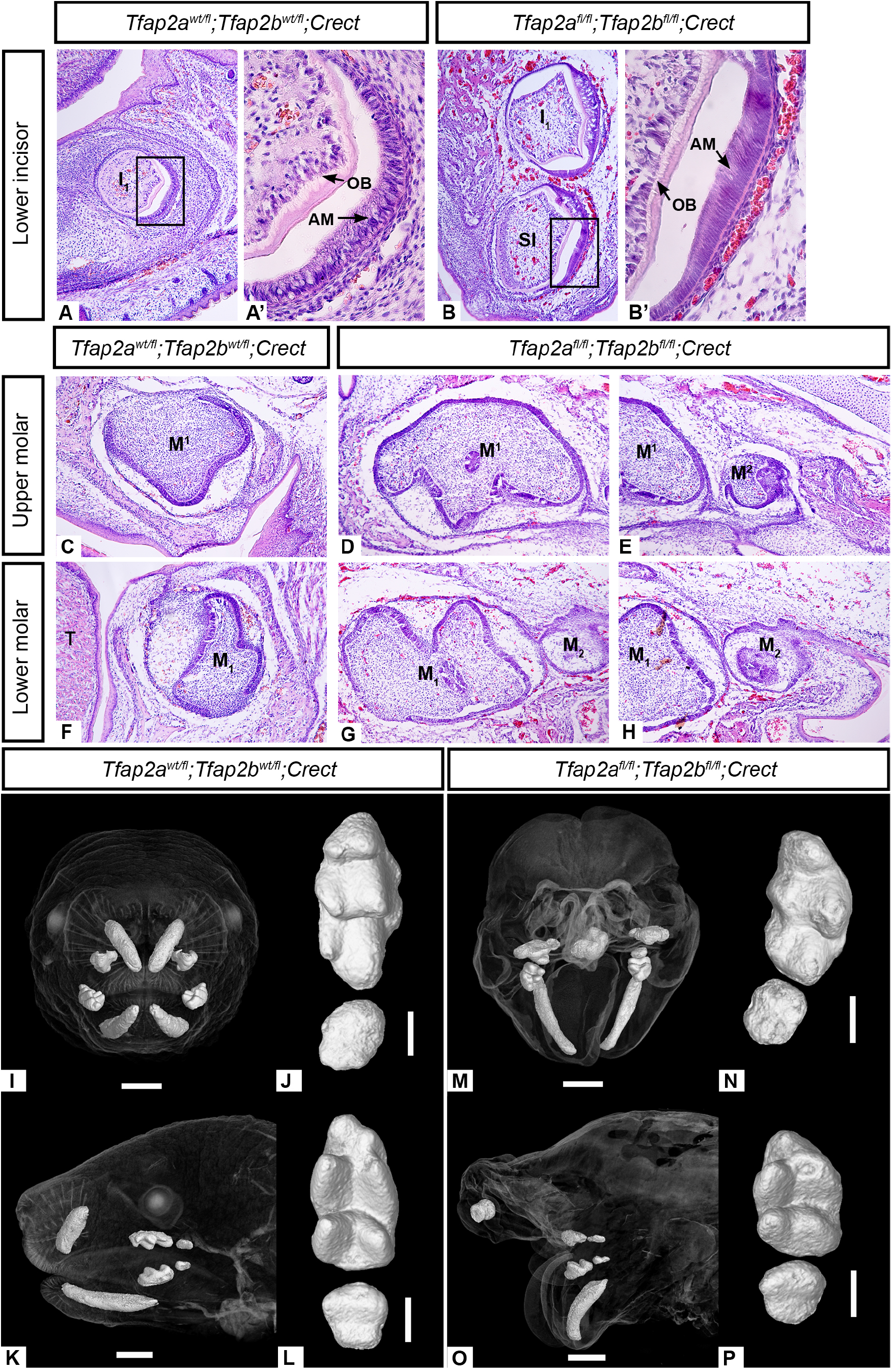
Duplicated lower incisors in E18.5 *Tfap2a^fl/fl^;Tfap^2bfl/fl^;Crect* mutant embryos contain ameloblasts and odontoblasts (B, B’) and mutants with only one incisor exhibit ventral curvature of this tooth (M, O). Hematoxylin and eosin staining of frontal cryosections showing that duplicated mutant incisors (B, B’) undergo cytodifferentiation at the bell stage similar to I_1_ and to control embryos (A, A’). In the mutant without duplicated incisors, a single ventrally curved lower incisor is present and two small upper incisors are present (M, O). In B and B’ the mutant mandibles exhibited ventral curvature (as seen in M, O), preventing the lower incisors and the upper and lower molars from being obtained in the same tissue sections, as in the controls. To ensure comparable planes of section with the controls, mutant hemi-mandibles were tilted backwards during embedding such that the sections through the lower incisors were taken through the anterior-most aspect of the mandible (true frontal plane). Note that only the right or left side of each frontal section is shown. Images taken of the embryos’ right side (A, A’, C, F, G, H) have been mirrored to match images taken of the left side. All histological images taken at 10X magnification. **3-D reconstructions from μCT data show that first and second molars develop in embryos lacking epithelial expression of** *Tfap2a* **and** *Tfap2b* **(D, E, G, H) and the cusp patterns on M^1-2^ (N) and M_1-2_ (P) look similar to the control (J: M^1-2^; L: M_1-2_).** Note that first molars are at the top and second molars are at the bottom. The mutant molars appear slightly shorter anterior-posteriorly than the control but all the main cusps are present. The apparent medial displacement of M^2^/_2_ relative to M^1^/_1_ in the mutant histological sections (E, H) is likely due to a slightly offset plane of section resulting from the cleft palate and mandible in the mutants. A-H taken at 10X; A’, B’ taken at 40X magnification. Scale bars are 1mm (I, K, M, O), and 0.3mm (J, L, N, P). AM: ameloblasts, OB: odontoblasts, SI: supernumerary incisor, T: tongue.

The face of the epithelium-specific mutants was highly dysmorphic (Van Otterloo et al., unpublished observations) which made it difficult to assess the development of the upper incisors. In some mutants, upper incisors were not observed at E18.5 (N=3/4 embryos) (Supplementary figure 6 G-I), but in one embryo we observed two small incisors though they were considerably shorter than those in the control (Figure 5 M, O; Supplementary figure 7 A-B; Supplementary table 2). In contrast to the incisors, the first and second upper and lower molars in the mutants were structurally similar to those of the controls, as assessed by histology (Figure 5 D, E, G, H). 3-D reconstructions from μCT scans and subsequent quantification of molar size (see Results 3.5), however, revealed that in mutants the molars were shorter along the mesiodistal axis and the cusps appeared less well defined than in the controls (Figure 5 N, P; Supplementary table 4).

We also examined mutant mice from a similar genetic cross in which one of the conditional alleles was null (*Tfap2a^flox/null^;Tfap2b^flox/null^;Crect*). In these mutants, we observed the same duplicated incisor phenotype in the lower incisors as in the first cross (Supplementary figure 8 D, E; Supplementary table 2), and one upper incisor was present (Supplementary figure 8 B). We also noted that the molars in these compound mutants were similar to the controls suggesting that the loss of one allele of each gene in the mesenchyme did not exacerbate the epithelium-specific mutant phenotype (Supplementary figure 8 F-K).

To determine how this supernumerary lower incisor develops, we also examined histological sections of cap stage (E14.5) teeth from *Tfap2a^flox/flox^;Tfap2b^flox/flox^;Crect* embryos. In hemi-mandibles from E14.5 control embryos (*Tfap2a^flox/wt^;Tfap2b^flox/wt^;Crect*), one cap stage lower incisor is present attached to the dental lamina (Figure 6 A), but in mutants, duplicated cap stage incisors were observed (N=2/3). In the mutants, I_1_ appeared tethered to the dorsal dental lamina (Figure 6 B, D) as in the controls, but the duplicated (ventral) incisor was tethered to the ventral surface epithelium, a region that does not normally have the characteristics of the dental lamina (Figure 6 C, E). In the mutant embryo lacking an extra cap-stage incisor, a bud emanating from the ventral surface epithelium was observed, indicating that this extra tooth was likely developmentally delayed compared to I_1_ (data not shown). Consistent with our observations at E18.5, upper incisors were not observed in *Tfap2a^flox/flox^;Tfap2b^flox/flox^;Crect* embryos at E14.5 (Supplementary figure 6 B-E) indicating that they failed to form prior to E14.5, at either the initiation or bud stage. Altogether, these results revealed a critical role for epithelial *Tfap2a* and *Tfap2b* function during tooth development.

**Figure 6.**
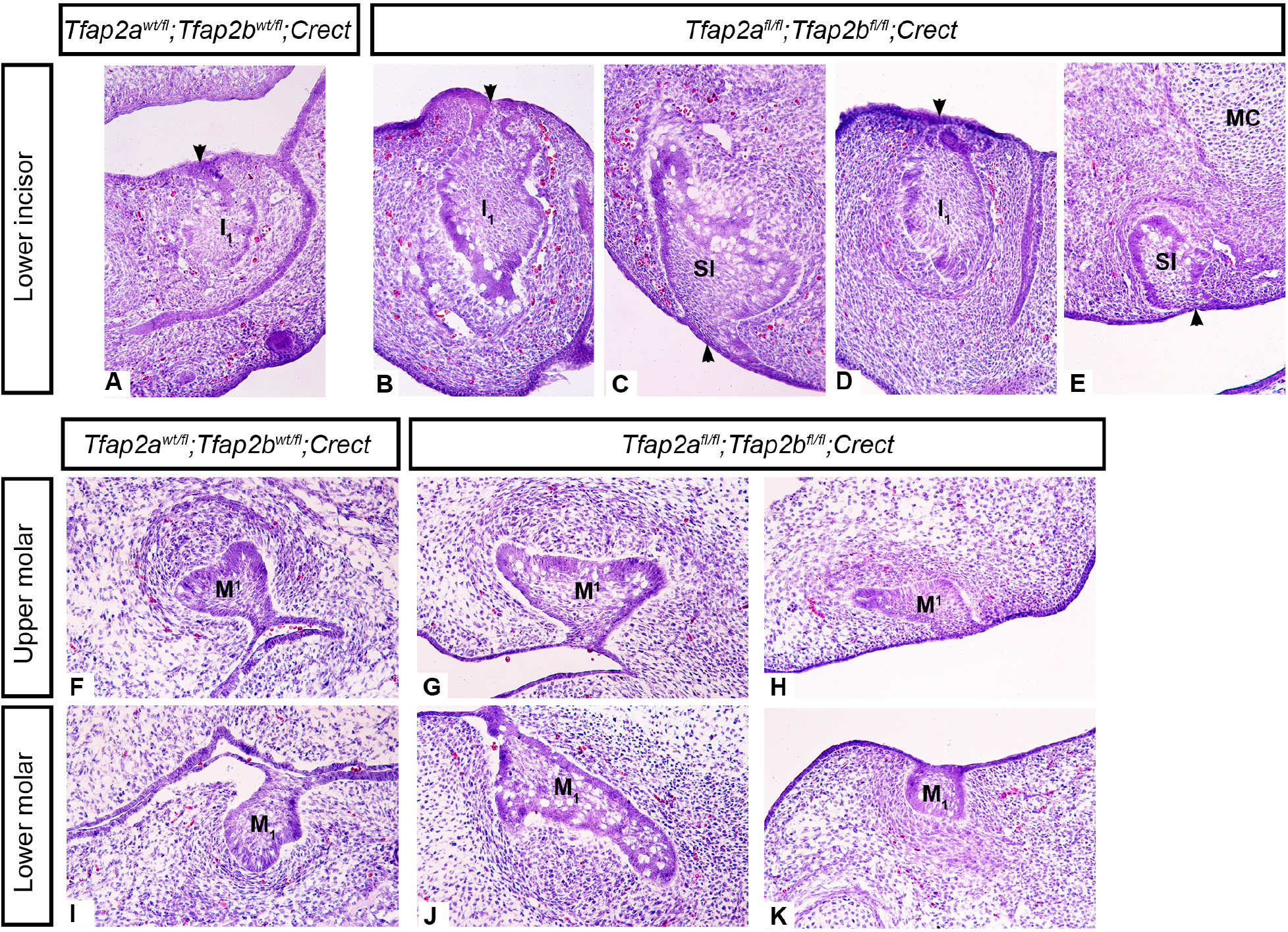
Supernumerary lower incisors are visible at E14.5 tethered to the ventral surface epithelium in *Tfap2a^fl/fl^;Tfap2b^fl/fl^;Crect* mutant embryos. Hematoxylin and eosin staining of frontal cryosections show that supernumerary incisors are ventral and slightly posterior to I_1_ at this stage. Representative sections of incisors from 2 individuals are shown (B+C, D+E). Note the attachment of the supernumerary incisors to the ventral epithelium (C, E, arrowheads) while I_1_ in the mutants (B, D) and controls (A) is attached to the dorsal dental lamina (arrowheads). Molars in mutant embryos vary slightly among individuals from early cap (H, K) to late cap stage (G, J). Aberrant appearance of molar teeth in frontal sections of *Tfap2* mutant embryos may be due to slight displacement of the molars along the medial-lateral axis relative to the plane of section as a result of cleft palate. SI: supernumerary incisor, MC: Meckel’s cartilage. Note that only the right or left side of each frontal section is shown. Images taken of the embryos’ right side (A, G, H, J, K) have been mirrored to match images taken of the left side. Images taken at 20X magnification.

### 3.4 Mesenchymal *Tfap2a* and *Tfap2β* is dispensable for tooth development, despite an interaction with ectodermal AP-2 function during jaw formation

Our findings described above show that *Tfap2a* and *Tfap2b* mRNA expression within the dental epithelium and/or oral epithelium serves an important function in proper tooth morphogenesis. Given that robust *Tfap2b* expression was detected in the dental mesenchyme (Figure 3), as well as some weaker domains, we predicted that AP-2 activity within the cranial neural crest-derived mesenchyme is also required for normal tooth development. To test this, we generated mice with conditional deletions of *Tfap2a* and *Tfap2b* in the neural crest-derived mesenchyme using the *Wnt1-Cre* allele (Danielian et al., 1998) and floxed or null alleles of *Tfap2a* and *Tfap2b* (Van Otterloo et al., 2018) (Supplementary figure 1 B, D).

Unexpectedly, and in contrast to the epithelial-specific loss of *Tfap2a* and *Tfap2b*, *Tfap2a^flox/flox^;Tfap2b^flox/flox^;Wnt1-Cre* embryos possessed the correct number of incisors and molars, and histology and μCT analysis confirmed that both classes of teeth are similar to controls at the cap E14.5 stage (Figure 7 A-F; Supplementary table 2) and at the bell (E18.5) stage (Figure 7 G-N). These results were replicated in a second cross with a null allele, *Tfap2a^flox/null^;Tfap2b^flox/null^;Wnt1-Cre,* (Supplementary figure 9 C, D, G, I; Supplementary table 2) and the lack of an apparent incisor phenotype was also noted in a previous study (Van Otterloo et al., 2018).

**Figure 7.**
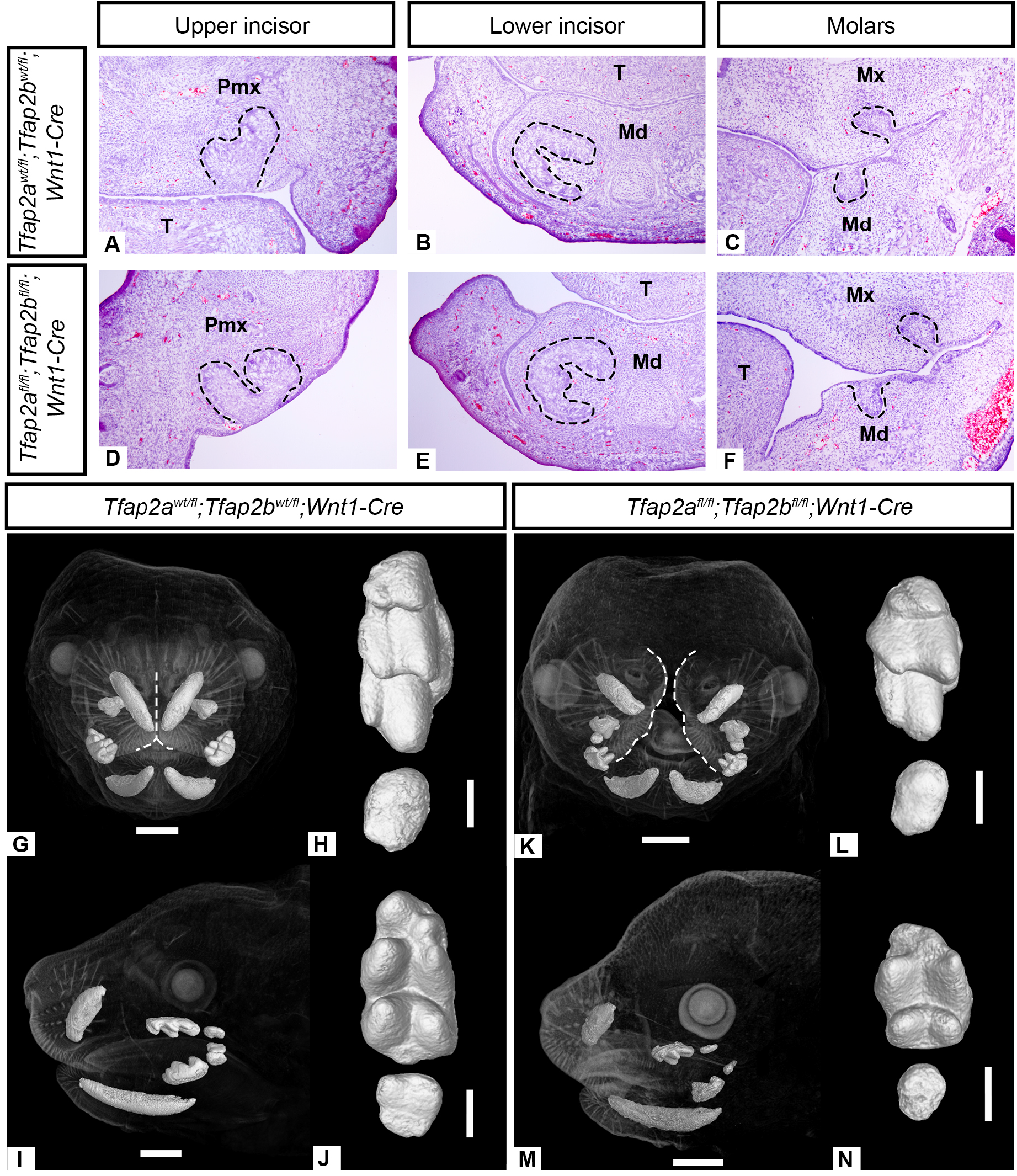
Incisors and molars in *Tfap2a^fl/fl^;Tfap2b^fl/fl^;Wnt1-Cre* embryos lacking *Tfap2a* and *Tfap2b* expression in the CNCC-derived mesenchyme lack major morphological defects based on histology (A-F) and μCT data (G-N). Hematoxylin and eosin stained cryosections in the frontal plane showing cap stage (E14.5) incisors (A, B, D, E) and molars (C, F) look similar in the mutant (D-F) and the control (A-C) embryos. Note that only the right or left side of each frontal section is shown. An image taken of the embryos’ right side (C) has been mirrored to match the corresponding images of the left side. Due to the clefted palate, the anterior frontal section (D) is angled on the medial aspect of the premaxilla. 3-D reconstructions of μCT data show that the upper molars (L: M^1-2^) and lower molars (N: M_1-2_) in the mutant appears shorter compared to the control (H: M^1-2^; J: M_1-2_) but all major cusps are present. Note the midface cleft in the mutant (K) outlined in white, compared to the control (G). A-F taken at 10X magnification. Scale bars are 1mm (G, I, K, M) and 0.3mm (H, J, L, N). Pmx: premaxilla, Mx: maxilla, Md: mandible, T: tongue.

A midface cleft and mandibular cleft were previously reported in mutant embryos in which *Tfap2a* and *Tfap2b* were deleted in the neural crest-derived mesenchyme and heterozygous in the ectoderm (*Tfap2a^flox/null^;Tfap2b^flox/null^;Wnt1-Cre*) (Van Otterloo et al., 2018) and we noted the same phenotype in this study (Supplementary figure 9 E). In contrast, in embryos with the mesenchyme-specific deletion of *Tfap2a* and *Tfap2b* and wild-type expression in the epithelium (*Tfap2a^flox/flox^;Tfap2b^flox/flox^;Wnt1-Cre*), we observed a midface cleft but no mandibular cleft (Figure 7 K).

### 3.5 Loss of *Tfap2a* and *Tfap2b* in either epithelium or mesenchyme leads to shorter molars

Histological analysis showed that loss of *Tfap2a* and *Tfap2b* in either the epithelium or the mesenchyme had little effect on the molar teeth; however, 3-D reconstructions of E18.5 embryos from both crosses revealed that the molars (M^1^/_1_) appeared shorter along the mesiodistal axis in the mutants compared to the controls (Figure 5 N, P; Figure 7 L, N). Quantification of the occlusal surface of the upper and lower first molars (ratio of molar length to molar width) from the 3-D reconstructions confirmed that for both crosses, each mutant embryo has shorter molars than the corresponding control embryo (Supplementary table 4). This phenotype was internally consistent among left and right sides (4 teeth/individual, although only one embryo per genotype was μCT-scanned). The molar ratios obtained from the control embryos, however, fall within the range observed in studies of wild populations of *Mus musculus* (Csanady and Mosansky, 2018; Wallace, 1968) and furthermore, the mutant M^1^/_1_, ratios fall outside of this range (Supplementary table 4). Given the similarity in molar ratios between the control embryos and wild mice, we used the standard deviation for M^1^ occlusal length from previously published measurements of wild adult *M. musculus* (Csanady and Mosansky, 2018) to estimate hypothetical distributions of occlusal length and width for our mutant and control mice and we performed a Wilcox-signed rank test to compare control versus mutant molar ratios from the estimated distributions. The results of this comparison were congruent with our initial observation that in both crosses, the mutant molars were shorter along the mesial-distal axis than those of the control embryos (p<2.2×10^−16^ alpha = 0.05; the same p-values were obtained for both crosses). We hypothesize that this difference in tooth length may be linked to the foreshortening of the snout in mutant embryos (Figure 5 O; Figure 7 M) noted here and in a previous study (Van Otterloo et al., 2018).

Collectively, these findings suggest that a complex interaction occurs between *Tfap2a* and *Tfap2b* function in the ectoderm (including oral and dental epithelia) and neural crest-derived mesenchyme during tooth and jaw development. These results also highlight that a neural crest-specific function for *Tfap2a* and *Tfap2b* is not necessary for embryonic tooth development.

## 4. Discussion

### 4.1 Temporal differences in duration of cranial neural crest migration to molars and incisors may influence pre-patterning of the dentition

The dental mesenchyme is a heterogeneous population of cells composed of both neural crest-derived cells and cells derived from the head mesoderm and the relative contribution of each of these cell populations to the dental mesenchyme changes during tooth development (Chai et al., 2000; Imai et al., 1996; Rothová et al., 2011). The origins of the cells that comprise the dental mesenchyme are of interest because shifts in the proportions of crest-derived or mesoderm-derived cells could have an effect on patterns of gene expression in the developing teeth.

CNCCs migrate into the branchial arches from approximately E7.5 to E9.0 in mice (Nichols, 1986, 1981; Theveneau and Mayor, 2012) and they fully invade the first branchial arch at the same time when the dentition is prepatterned into future incisor and molar regions (E9.5-E10.5) (Lumsden, 1988). Previous cell fate mapping using DiI labeling in rats revealed that early migrating CNCCs populate anterior regions of the face and jaw, whereas late-migrating CNCCs populate more posterior regions (Imai et al., 1996). To build upon the results of this earlier study, we explicitly tested whether there are differences in the neural crest contribution to developing incisors and molars. We performed a time-course lineage tracing experiment in which CNCCs were labeled at three different time points during migration. Our results show that CNCCs migrate to the future incisor region and molar regions at ~E7-E8. Late-migrating (E8.5) CNCCs, however, predominantly populate the molar mesenchyme, while relatively few of these cells invade the incisor mesenchyme. Labeling CNCCs for the majority of the duration of their migration produced equivalent results to the E6.5 or E7.5 single-day labeling, further demonstrating that the temporal difference in neural crest contribution between incisors and molars is limited to late-migrating crest cells.

Earlier experiments in mice established that tissue grafted from E10 frontonasal prominences gave rise exclusively to incisors, whereas mesenchyme from the maxillary prominences formed molars (Lumsden and Buchanan, 1986). Additionally, CNCCs that migrate into the premaxilla and maxilla originate from two different locations in the neural ectoderm; the premaxilla and incisor mesenchyme are populated by crest cells derived from the forebrain and midbrain while the maxilla and molar mesenchyme form from crest cells derived from the midbrain and hindbrain (rhombomeres 1 and 2) (Jiang et al., 2002).

Cranial neural crest-derived mesenchyme by itself is insufficient for tooth formation which requires inductive cues from the dental epithelium and likewise, the epithelium alone is also unable to produce teeth (Mina and Kollar 1987, Lumsden, 1987, 1988). When considered in the context of earlier work, our data suggest that the differences in the duration of neural crest cell migration into the incisor and molar regions may contribute to the anterior-posterior patterning of the dentition. This difference in duration of migration could affect the molecular response of the mesenchyme to initial inductive cues from the dental epithelium, essentially helping to facilitate differential gene expression in anterior versus posterior dental mesenchyme.

### 4.2 Expression patterns of *Tfap2a*, *Tfap2b* differ between incisors and molars during dental development

To investigate the potential molecular differences in CNCCs in incisors and molars, we compared expression patterns of two AP-2 transcription factors which are expressed in neural crest cells (Mitchell et al., 1991; Moser et al., 1997). Spatiotemporal gene expression analyses revealed that expression domains of *Tfap2a* and *Tfap2b* differ among incisor, molar, oral epithelium, and surface epithelial tissues. Differences in expression were observed between incisors and molars as tooth development proceeds from bud, to cap, to bell stage. At the bud stage (E12.5 and E13.5) and cap stages (E14.5) both genes were expressed in the surface epithelium, while in the upper and lower incisors *Tfap2a* was expressed prominently in the dental epithelium compared with weaker expression of *Tfap2b.* In the incisor mesenchyme at the late bud and cap stages, only *Tfap2a* was observed. Epithelial and mesenchymal *Tfap2a* expression persisted into the bell stage in the incisors and *Tfap2b* expression became restricted to the incisor inner enamel epithelium, including ameloblasts.

In the molar epithelium, *Tfap2a* was detected but *Tfap2b* was essentially absent at the bud and cap stages. Conversely, in the molar mesenchyme from the bud through cap stages, *Tfap2a* and *Tfap2b* were both expressed, albeit weakly for *Tfap2a* and more robustly for *Tfap2b*, consistent with previous reports (Tanasubsinn et al., 2017; Uchibe et al., 2012). By the bell stage *Tfap2a* was prominently expressed in the molar inner enamel epithelium while *Tfap2b* was restricted to molar mesenchyme in the outer region of the tooth germ. Restricted expression of *Tfap2a* in the molar inner enamel epithelium suggests it may be associated with proliferation or folding of the molar inner enamel epithelium, which is thought to be regulated by enamel knots that facilitate the development of multiple cusps on the molar surface (Cho et al., 2007; Matalova et al., 2005; Thesleff et al., 2001).

Previous work showed that YEATS4 increases the transcriptional activity of TFAP2B (Ding et al., 2006) while KCTD1 inhibits the transcriptional activity of TFAP2A or TFAP2B (Ding et al., 2009). We detected overlapping expression domains between *Tfap2a*, *Kctd1,* and *Yeats4* in cap stage dental epithelium in incisors and molars which suggests that *Kctd1* and *Yeats4* may interact directly with *Tfap2a* to modulate its expression in developing teeth, though this prediction remains to be tested. Additionally, at the early bell stage, *Yeats4* and *Tfap2b* were both expressed in incisor ameloblasts. These findings highlight regions of both co-expression and divergent expression domains of *Tfap2a* and *Tfap2b* and their regulators during development of incisors and molars.

### 4.3 Cooperative functional roles for TFAP2A and TFAP2B in the craniodental ectoderm during tooth formation

It has been well established that TFAP2A and TFAP2B are able to form heterodimers (Ding et al., 2009; Williams and Tjian, 1991), can bind the same DNA consensus sequences (Williams and Tjian, 1991), and are capable of functioning redundantly in tissues in which they are co-expressed (Hoffman et al., 2007; Li and Cornell, 2007; Rothstein and Simoes-Costa, 2020; Seberg et al., 2017; Van Otterloo et al., 2018; Wang et al., 2008). We showed that TFAP2A and TFAP2B cooperatively function within the craniodental ectoderm, including the dental epithelium, to regulate incisor development. Conditional deletion of these genes specifically within the craniodental ectoderm led to duplication or ventral curvature of lower incisors and loss or reduction of the upper incisors. It remains to be determined how epithelial AP-2 function may influence incisor development and the spatiotemporal requirements of this function.

Two potential explanations for the lower incisor duplication in mutants involve a role for AP-2 in dorsoventral patterning. One possibility is that early in dental development, the odontogenic region is expanded ventrally, allowing the initiation of an ectopic incisor within the non-odontogenic aboral domain of the mandible (Figure 8). Alternatively, loss of AP-2 function within the surface ectoderm could indirectly affect incisor number by shifting the dorsoventral axis identity within the mandible such that the ventral (aboral) region takes on a more dorsal-like identity, thereby permitting the formation of an ectopic tooth.

**Figure 8.**
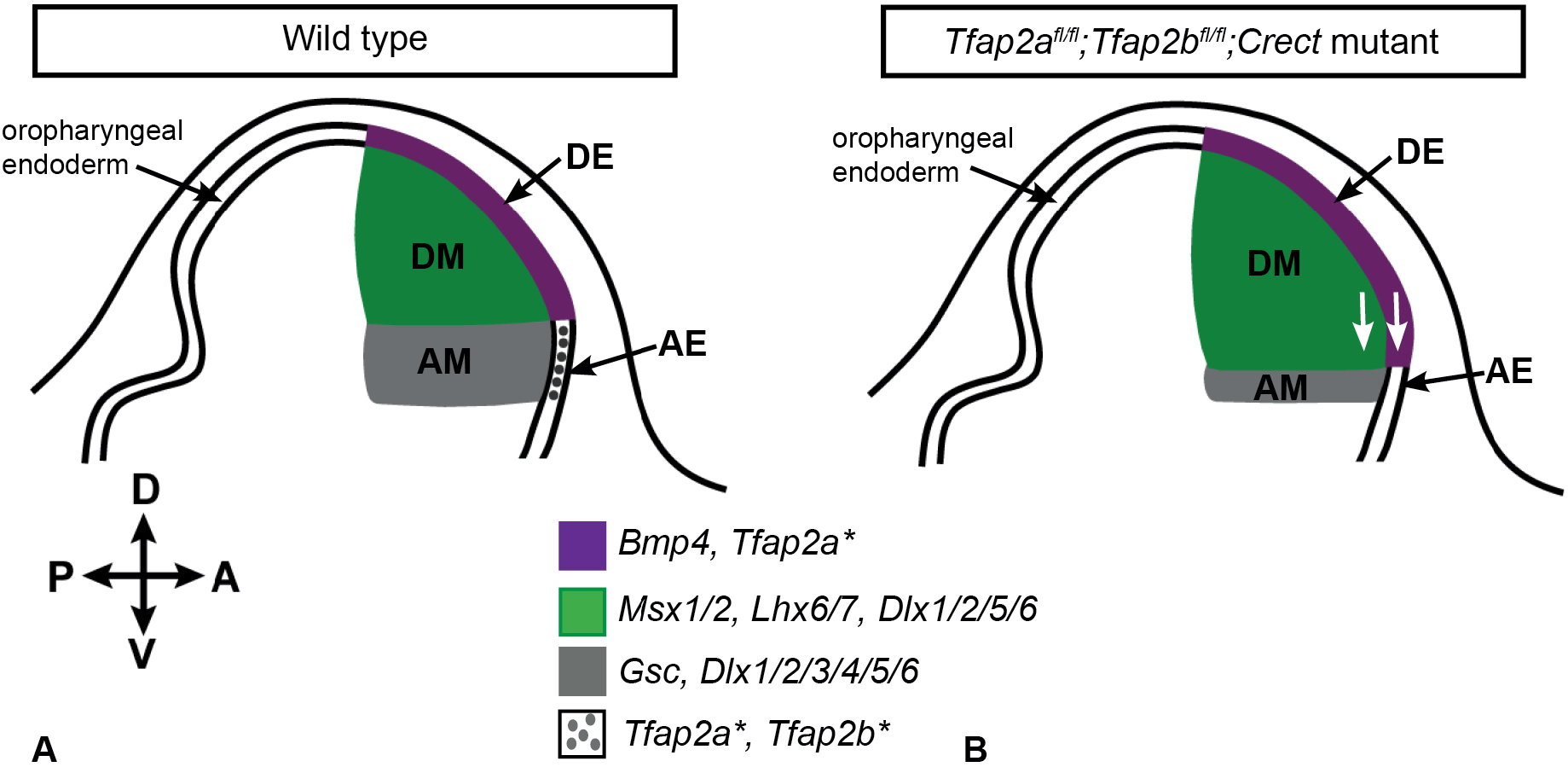
Model for development of duplicated lower incisors in mice lacking *Tfap2a* and *Tfap2b* expression in the epithelium. Schematic drawing of a sagittal section through a mouse mandible at E10.5, showing the incisor (medial) region for the wild type (A) and the mutant (B). In this model, loss of function of *Tfap2a* and *Tfap2b* in the epithelium leads to dorsoventral mis-patterning in the anterior aspect of the mandible. We hypothesize that this perturbation of the dorsoventral axis results in ventral expansion of the odontogenic domain into the aboral epithelium and mesenchyme (white arrows), perhaps via expansion of *Bmp4, Lhx6/7, Msx1/2* expression domains. In our model, ventral expansion of the odontogenic domain could result in the initiation of ectopic tooth buds that were limited to the anterior (incisor) region of the developing mandible, thereby leaving the molar buds unaffected, as observed in the epithelium-specific *Tfap2a/Tfap2b* mutants. Asterisks indicate genes that have been deleted in the mutant embryos. DE: dental epithelium, DM: dental mesenchyme, AE: aboral (surface) epithelium, AM: aboral mesenchyme.

Given that in our ectoderm-specific line (*Crect*), *Cre*-recombinase is expressed in both epithelial (including oral and dental epithelia) and surface ectoderm tissues, it is difficult to distinguish between these models. A complete transformation of a mandibular axis would likely also affect the molars, unless the transformation was isolated to anterior/distal elements of the mandible, therefore this seems improbable in these mutants. Future studies utilizing an alternate *Cre*-recombinase line that specifically targets the dental epithelium and/or expression profiling would be needed to further distinguish between these two hypotheses.

Interestingly, in the E14.5 mutant embryos in which a supernumerary cap stage lower incisor was observed (*Tfap2a^flox/flox^;Tfap2b^flox/flox^;Crect*), it appeared to be connected to the ventral surface epithelium (Figure 6 C, E), as if the surface epithelium were an ectopic dental lamina. This suggests that the ectopic tooth initiated in the ventral epithelium, but examination of earlier-stage embryos (*e.g.,* ~E11.5) would be needed to test this hypothesis. Ventral curvature of I_1_ in the mutants lacking duplicated incisors is also suggestive of dorsoventral mis-patterning, however, this may be secondary to changes in the shape of the mandible which also curves ventrally compared to the control (Figure 5 M, O) (Van Otterloo et al., unpublished observations). Properly patterned molars in the mutants demonstrates that the anterior teeth (incisors) are acutely affected by the loss of expression of *Tfap2a* and *Tfap2b*.

Altogether, the incisor duplication or ventral curvature observed in the mutant embryos suggest that *Tfap2a* and *Tfap2b* expression in the epithelium may be important for establishing or maintaining dorsoventral polarity within the anterior aspect of the mandible. Specifically, these results imply that an additional ventral epithelial region with the capacity to fully execute the odontogenic program is established along with the dental lamina in the oral cavity and that the non-dental mesenchyme is also competent to respond to initiation cues emanating from the supernumerary dental epithelium. Lack of additional ectopic tooth germs in this region indicates that the duplicated tooth likely forms from a spatially restricted region, similar to the native dental lamina.

Despite robust *Tfap2a* and *Tfap2b* expression within the molar tooth germs, we did not detect major defects, aside from increased tooth length, within this tooth class following loss of *Tfap2a/Tfap2b* from either the epithelial or mesenchymal tissues. There are several potential explanations as to why the molars were largely unaffected. First, it is possible that despite prominent expression, AP-2 function is not required for molar development. Second, additional *Tfap2* family members, namely *Tfap2c*, may compensate for the loss of *Tfap2a* and *Tfap2b*. Expression of *Tfap2c* was previously identified within the oral epithelium and dental mesenchyme (Chazaud et al., 1996) and it would be interesting to test whether compound loss of *Tfap2a, Tfap2b,* and *Tfap2c* would perturb molar development. A third related possibility is that loss of *Tfap2a/Tfap2b* in either the epithelium or mesenchyme alone is insufficient to disrupt molar development due to compensation from expression within the alternate tissue. In this scenario, elimination of *Tfap2* from both epithelium and mesenchyme simultaneously would be required to significantly disrupt molar development. Mice that are homozygous null for *Tfap2a*, however, are so severely affected that they lack a mouth and other ventral craniofacial structures (Zhang et al., 1996), and therefore a temporally inducible deletion would likely be needed to address this.

### 4.4 Phenotypic variability in incisors following loss of epithelial TFAP2A/TFAP2B function both within and between genetic crosses

Supernumerary lower incisors are present in both *Tfap2a^flox/flox^;Tfap2b^flox/flox^;Crect* embryos (*i.e., Tfap2a/Tfap2b* are “wild type” outside the *Cre*-positive domain) and *Tfap2a^flox/null^;Tfap2b^flox/null^;Crect* embryos (*i.e., Tfap2a/Tfap2b* are *heterozygous* outside of the *Cre*-positive domain). In both crosses, however, we also observed embryos in which the incisor exhibited ventral curvature but was not duplicated. Variability within and between crosses was also noted for the upper incisors, which were either absent, reduced from two incisors to one, or reduced in size. Larger sample sizes would be needed to better characterize the variation in the upper incisor phenotype in these mutants.

### 4.5 Potential molecular mediators of dental defects in *Tfap2a/Tfap2b* ectodermal mutants

If an odontogenic region has been expanded or duplicated in *Tfap2a/Tfap2b* ectodermal mutant embryos, what molecular signals are responsible? It has been well-established in mice that the dental epithelium is patterned into two distinct regions, an anterior incisor region and a more posterior molar region, and the inductive molecular signals from each region in the epithelium elicit different molecular responses from the dental mesenchyme (Chen et al., 1996; Neubüser et al., 1997; Tucker et al., 1998b; Xu et al., 2019). The proximal-distal axis of the dentition is established by E10.5 within the presumptive dental epithelium, via expression of *Bmp4* anteriorly in the future incisor region and *Fgf8* posteriorly in the future molar region (Neubüser et al., 1997; Xu et al., 2019). In response to epithelial expression of *Bmp4, Msx1/2* are expressed in the underlying incisor mesenchyme (Chen et al., 1996; Tucker et al., 1998b) and epithelial expression of *Fgf8* induces *Pax9* expression in the molar mesenchyme (Neubüser et al., 1997).

Demarcation of the oral-aboral (dorsal-ventral) axis of the mandible also occurs early in development primarily via complementary expression of homeobox genes, including, *Goosecoid* (*Gsc*), which is limited to the non-dental mesenchyme, and *LIM homeobox 6* and *7* (*Lhx6* and *7*), which are expressed in overlapping domains throughout the presumptive dental mesenchyme (Grigoriou et al., 1998). Previous work demonstrated that at E10 the aboral (ventral) mesenchyme is competent to form teeth when cultured with exogenous *Fgf8* which induces expression of *Lhx6/7* and represses *Gsc*, but that this ability is lost by E11 when the aboral mesenchyme is no longer competent to express *Lhx6/7* (Tucker et al., 1999).

Given that the incisors are uniquely affected in the mutant embryos that lack *Tfap2a* and *Tfap2b* expression in the epithelium and that the duplicated incisor appears to be developing along the same timeline as I_1_ we predict that a ventral dental lamina is patterned and that a supernumerary tooth germ is initiated at the same time as I_1_ (E10-11) when the mesenchyme, both oral and aboral, is competent to respond to initiating cues from the dental epithelium. At E10.5, *Tfap2a* is expressed in the crest-derived mesenchyme of the first branchial arch while both *Tfap2a* and *Tfap2b* are expressed in the epithelium surrounding the first branchial arch (Zhang and Williams, 2003; Zhao et al., 2011). Our expression data show that *Tfap2a* and *Tfap2b* are also expressed in the dental and mandibular epithelia but only *Tfap2a* is expressed in the incisor mesenchyme at the early bud stage (E12.5). The transient inductive capacity of the epithelium and competence of the branchial arch mesenchyme (oral and aboral) to respond at E10-11, may explain why the loss of *Tfap2a* and *Tfap2b* expression in the epithelium has a much larger effect on early patterning of the dentition compared to the deletion of *Tfap2a* and *Tfap2b* in the neural-crest derived mesenchyme.

We hypothesize that expansion and/or upregulation of *Bmp4* signaling in the epithelium at the time when the oral-aboral axis of the first branchial arch is established (~E10-10.5) could result in a ventral expansion of the anterior odontogenic domain (Figure 8). Previous work showed that in *Tfap2b*-null mice, *Bmp4* is upregulated and slightly expanded in the distal limb buds at E10.5-11.5 and that *Bmp4* promoter activity is negatively regulated by both *Tfap2a* and *Tfap2b* (Zhao et al., 2011). If loss of epithelial expression of *Tfap2a* and *Tfap2b* also results in loss of negative regulation of *Bmp4* in the face as in the limbs, this could result in upregulation and/or expansion of *Bmp4* expression in the future incisor region at ~E10.5. Upregulation of *Bmp4* at E10.5 could, in turn, result in expansion/upregulation of downstream genes (e.g., *Msx1*) in the dental mesenchyme. *Msx1* normally becomes restricted to the dorsal (odontogenic) mesenchyme within the mandible by ~E11 (Tucker et al., 1998a). Importantly in this model, upregulation/expansion of *Bmp4* and, subsequently, *Msx1* expression at ~E10.5 could hypothetically lead to an additional incisor domain without affecting patterning more posteriorly in the molar region, demarcated by epithelial *Fgf8,* therefore, this could result in the phenotype observed in embryos from both crosses lacking epithelial *Tfap2a* and *Tfap2b* expression (*Tfap2a^flox/flox^;Tfap2b^flox/flox^;Crect* and *Tfap2a ^flox/null^;Tfap2b ^flox/null^;Crect*).

Additionally, *Dlx* genes are crucial for proximal-distal patterning the jaw and dentition (Depew et al., 1999; McCollum and Sharpe, 2002; Qiu et al., 1997; Zhao et al., 2000), and *Dlx2, 5,* and *6* are expressed in the oral mesenchyme of the mandible (Zhao et al., 2000). Given that all six *Dlx* genes are downregulated in mice lacking mesenchymal expression of *Tfap2a* and *Tfap2b* (Van Otterloo et al., 2018), and it seems likely that expression of these genes, particularly *Dlx2, 5,* and *6*, may also be affected in the *Tfap2a* and *Tfap2b* ectoderm-specific (*Crect*) mutants.

## Acknowledgements

We thank Dr. Brooke Armfield for her assistance with experimental design, mouse breeding, and for teaching EDW to perform various assays, including *in situ* hybridization. We thank Dr. Christine Larkins and Dr. Kelsey Lewis for assistance and advice with the neural crest lineage tracing experiment, Alyssa Mangino for assistance with sectioning and *in situ* hybridization, Blake Hauer for assistance with sectioning, and Emily Merton for technical support. We also acknowledge Dr. Gary Scheiffle and Dr. Edward Stanley at the University of Florida Nanoscale Research Facility for their assistance with μCT scanning. Finally, we thank all members of the Cohn laboratory for valuable insights and critical discussion of this work.

## Data Availability

The μCT data from this study will be freely available to the public on FaceBase3 (https://www.facebase.org/).

