## Supplementary Materials for "Anomalous incisor morphology indicates tissue-specific roles for *Tfap2a* and *Tfap2b* in tooth development"

### 5. Supplementary Methods

#### 5.1 Lineage tracing of *Sox10*-positive cranial neural crest cells

***CNCC lineage tracing:*** In *Sox10-iCre/ER$^{T2}$* embryos, *Cre*-mediated recombination is driven by the *Sox10* promoter and is induced by tamoxifen. In the presence of tamoxifen, the iCRE/ER$^{T2}$ fusion protein binds tamoxifen and translocates to the nucleus where it recombines with the *R26R$^{mTmG}$* transgene, resulting in excision of the tdTomato sequence and expression of EGFP in *Sox10*-expressing CNCCs. Tamoxifen (Sigma-Aldrich T5648) was mixed with filtered corn oil (40 mg tamoxifen per 1 ml corn oil) and incubated at 37°C for 1.5-2 hours prior to administration. For both lineage tracing experiments, we administered one dose of tamoxifen (200 mg tamoxifen/kg mouse weight) to pregnant female mice by oral gavage either on one single day (experiment 1: E6.5, E7.5, or E8.5) or on three consecutive days (experiment 2: E6.5-E8.5).

***Embryo processing:*** E13.5 embryos were dissected, fixed briefly in 4% PFA for 1 hour on ice in the dark, rinsed in PBS, and screened for EGFP and tdTomato fluorescence. EGFP-positive (*Cre*-positive) and tdTomato-positive embryos were prepared for cryosectioning as described above (Methods 2.3). Results reported are based on a minimum of 3 embryos for each time point within each of these experiments. After sectioning, tissue sections were briefly fixed in 4% PFA for 10 minutes, rinsed twice in PBS and mounted on slides using Fluoromount-G fluorescent mounting medium (Southern Biotech) and covered with glass coverslips (Thermo Fisher Scientific). Membrane fluorescence for EGFP and tdTomato was visualized in frontal cryosections using a Zeiss LSM 710 confocal microscope.

***Genotyping:*** To genotype embryos, cells were lysed in sodium hydroxide (25mM) at 95°C for 1 hour, neutralized with tris hydrochloride (40mM), and diluted to ~5ng/µl. PCR for both *R26R$^{mTmG}$* and *Sox10-iCre/ER$^{T2}$* alleles was performed using primers specific to each transgene (Supplementary table 1) using the following conditions: 95°C for 3 minutes, 95°C for 30 seconds, 64°C for 30 seconds, 72°C for 1 minute for 35 cycles.

#### 5.2 Tissue-specific conditional deletion of *Tfap2a* and *Tfap2b*

In double mutant embryos generated from the first cross (*i.e., Tfap2a*$^{flox/flox}$*;Tfap2b*$^{flox/flox}$*;Cre+*), deletion of *Tfap2a* and *Tfap2b* was strictly isolated to the tissue of interest (*i.e.,* the *Cre*-positive tissue), whereas *Cre*-negative tissues retained *Tfap2a* and *Tfap2b* function and were essentially 'wild-type' (Supplementary figure 1 A, B). In the second cross, however, double mutant embryos (*i.e., Tfap2a*$^{null/flox}$*;Tfap2b*$^{null/flox}$*;Cre+*) lacked both copies of *Tfap2a* and *Tfap2b* in *Cre*-expressing cells but remained heterozygous for *Tfap2a* and *Tfap2b* in the rest of the embryo (Supplementary figure 1 C, D). Embryos that were heterozygous for both *Tfap2a* and *Tfap2b* conditional alleles (*i.e., Tfap2a*$^{wt/flox}$*;Tfap2b*$^{wt/flox}$*;Cre+*) were used as controls. All *Tfap2* mouse lines were initially maintained on a mixed 129/Black Swiss background (Brewer et al., 2004) but they have since been outcrossed to Black Swiss (strain code 492, Charles River).

For the experiments involving *Tfap2* mutant embryos, DNA was extracted by placing the isolated tissue in DirectPCR Lysis Reagent (Viagen Biotech) plus 10 µg/ml proteinase K (Roche), incubated overnight at 65°C, followed by heat inactivation at 85°C for 45 min. The lysed product was then used directly in a PCR reaction using the Qiagen DNA polymerase kit, including the optional Q Buffer solution (Qiagen).

#### 5.3 Cloning of RNA probes for *in situ* hybridization

***Primer and probe design:*** Oligonucleotide primers (Supplementary table 3) were designed in Geneious (v6.1.8 or 10.0.9, Biomatters, Ltd) such that the resulting amplicon (DNA template for the RNA probe) was from a single exon. To ensure that the probes were specific to the genes of interest, we performed BLAST (Basic Local Alignment Search Tool) searches of the NCBI database (http://blast.ncbi.nlm.nih.gov/).

***Cloning:*** PCR products were visualized using gel electrophoresis. The PGem-T Easy vector (Promega) or the Strataclone vector (Strataclone) were used to clone the target DNA fragments. These vectors contain M13 forward and reverse priming sites and the following priming sites and polymerase start sites that were utilized in downstream steps in this protocol:

T7 (both vectors), T3 (Strataclone vector), and Sp6 (PGem-T Easy vector). Amplicons were ligated into the vector and cloned using NEB Turbo competent cells (New England Biolabs) grown on plates containing Luria-Bertani (LB) broth medium with ampicillin and X-gal was used for blue/white screening of bacterial colonies. Transformed bacterial colonies were grown with agitation (7 hours at 300 RPM or 12 hours at 225 RPM) in Terrific Broth media containing ampicillin. Cultures were purified using the GenElute Miniprep kit (Sigma Aldrich) and linear inserts were obtained using PCR with M13 forward and reverse primers. PCR products were purified using the Promega PCR and Gel Clean-Up kit (Promega) and gel electrophoresis was used to verify that the cloned fragments were the correct size. We determined the orientation of the target sequence within the plasmid and validated the identity of each target gene sequence by Sanger sequencing. “Antisense” and “sense” DIG-labeled RNA probes were synthesized in a transcription reaction using the appropriate polymerase (T7, T3, or Sp6 (Promega)), purified using the RNA Mini Quick Spin Columns (Roche) and the concentration (ng/µl) of each probe was measured using a Nanodrop spectrophotometer (Thermo Fisher Scientific).

#### 5.4 *In situ* hybridization on cryosections

Steps were performed at room temperature unless noted. Cryosections were thawed briefly, fixed in 4% PFA with PBS (10 min), washed twice in DEPC-treated PBS + 0.1% Triton-X 100 (DEPC-PBST) and treated with proteinase K (5µg PK/ml PBST, 5 min). Tissue was then washed with DEPC-PBST and fixed again in 4% PFA (5 min), followed by an acetylation wash to increase the specificity of the ISH signal (Braissant and Wahli, 1998). Sections were incubated in warm pre-hybridization solution at 65°C (1-2 hours). DIG-labeled “antisense” and “sense” RNA probes were diluted to concentrations ranging from 0.2-0.5 µg/µl in warm prehybridization solution and 0.2-0.5ml of this solution was immediately applied to each slide. Slides were incubated at 65°C overnight in a humid hybridization chamber.

Slides were rinsed in a series of saline-sodium citrate (SSC) solutions of decreasing stringency at 65°C: 5X SSC, 2X SSC + formamide + 0.1% CHAPS, and 0.2X SSC + 0.1% CHAPS. Slides were rinsed in KTBT solution (50mM TrisHCl pH 7.5, 150mM NaCl, 10mM KCl, 1% Triton X-100), blocked with 20% goat serum in KTBT (2 hrs at 4°C), and incubated overnight at 4°C in anti-DIG-AP antibody (Roche) diluted in blocking buffer (1:3,000). Multiple KTBT washes (30-45 min each) were performed to remove non-specifically bound antibody, followed by washes with NTMT solution (100mM Tris HCl pH 9.5, 100mM NaCl, 50mM $MgCl_2$, 0.1% Triton X-100). Sections were placed in NBT-BCIP coloring solution for multiple days. When the coloring was complete, the slides were rinsed in KTBT, fixed in 4% PFA, mounted with Dako glycergel aqueous mounting medium (Agilent Technologies) and covered with glass coverslips (Thermo Fisher Scientific). Tissue sections were photographed on a Leica microscope using Kohler illumination and any subsequent adjustments made to brightness/contrast were performed in Adobe Photoshop CS6 (v13.0.6) on the entire image.

**5.5 Embryo processing for micro-CT scanning**

E18.5 embryos for micro-CT scanning were fixed in 4% PFA, rinsed in PBS, and stained in the dark with a contrast-enhancing agent, Lugol's iodine solution (1.25% $I_2$, 2.5% KI with DEPC water) overnight at room temperature. Excess iodine solution was removed with a Kim Wipe and embryos were placed in a small plastic capsule to prevent desiccation during scanning. After scanning, embryos were rinsed in 70% ethanol and then PBS to remove the iodine stain and were subsequently processed for histological analysis as described above (see Methods 2.4). Lugol's solution causes some degree of tissue shrinkage (Metscher, 2009), but we expect that this shrinkage was similar across same-stage embryos because the duration in Lugol's was the same for all embryos.

### Supplementary Figures and Tables

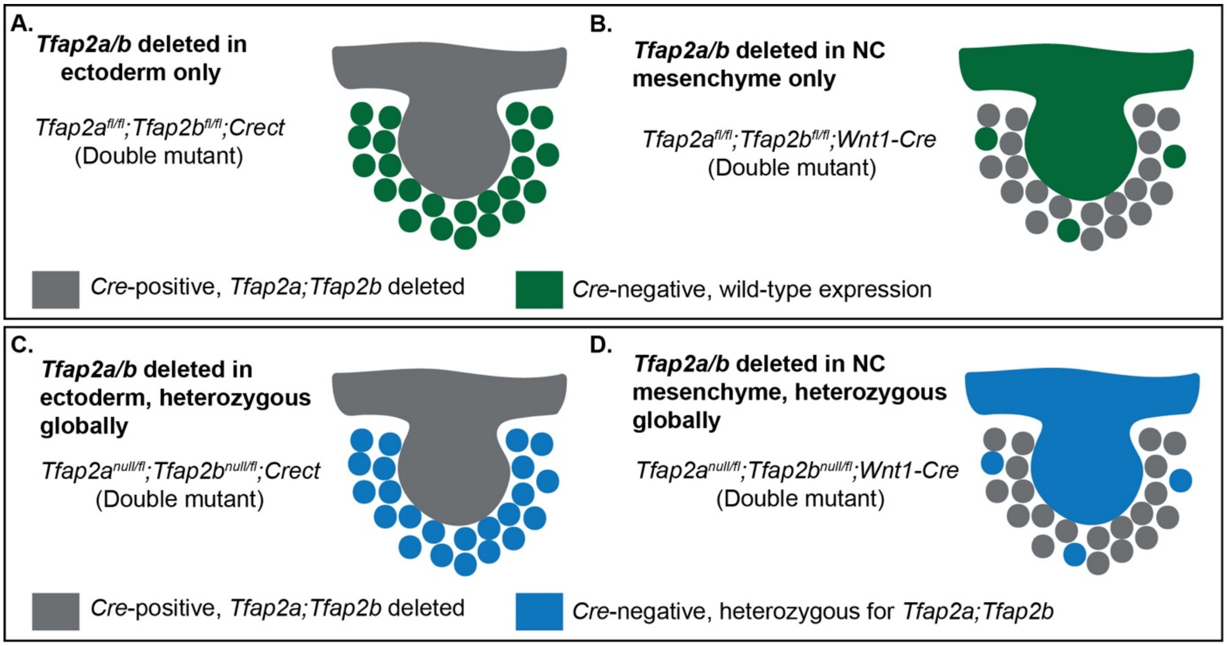


**Supplementary Figure 1. Schematic illustrating *Tfap2a* and *Tfap2b* double conditional mutant genotypes depicting where *Tfap2a* and *Tfap2b* are deleted in the developing teeth in each tissue-specific cross.** *Tfap2a* and *Tfap2b* are deleted in the ectoderm in *Crect* crosses (A, C) while in the *Wnt1-Cre* crosses, both genes are deleted in the neural crest-derived mesenchyme. Note that mesoderm-derived mesenchyme makes a minor contribution to the dental mesenchyme, shown here with a few colored mesenchyme cells that are *Wnt1-Cre* negative (B, D). In crosses utilizing conditional alleles for *Tfap2a* and *Tfap2b* (A, B), the *Cre*-negative tissues are “wild type” (green shading). In contrast, crosses that include *Tfap2a* and *Tfap2b* null alleles (C, D), *Cre*-negative tissues are heterozygous for *Tfap2a* and *Tfap2b* (blue shading).

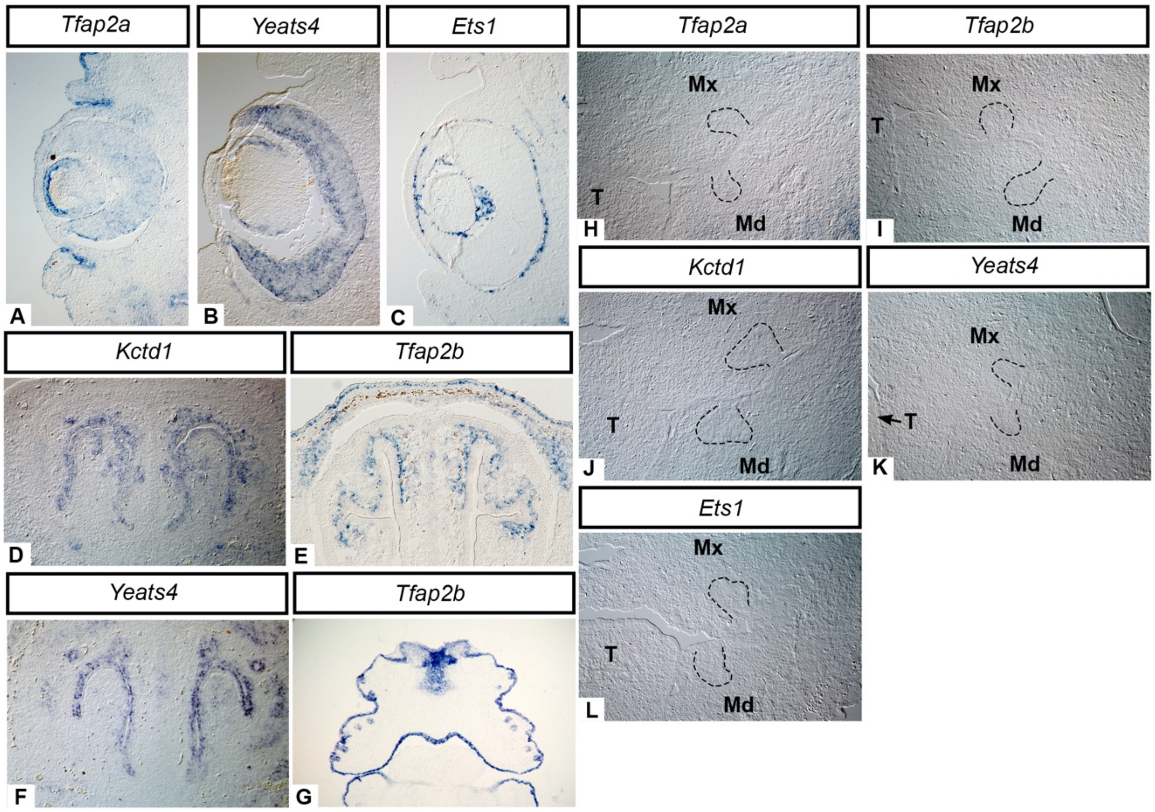


**Supplementary Figure 2. Positive and negative controls for *in situ* hybridization for *Tfap2a, Tfap2b, Yeats4, Kctd1,* and *Ets1*.** Cryosections in the frontal plane showing that at E14.5 *Tfap2a, Yeats4, and Ets1* were detected in the eye (A-C), while *Kctd1,* and *Yeats4* were detected in the nasal epithelium (D, F) and *Tfap2b* was expressed in the nasal mesenchyme (E). *Tfap2b* was also expressed in the facial (surface) epithelium at E14.5 (G). Transcripts were not detected with the "sense" mRNA probes in frontal cryosections from molar tooth germs at E14.5 *Tfap2a* (H), *Kctd1* (J) *Ets1*, (L) or E13.5 *Tfap2b* (I), *Yeats4* (K). The molar epithelium is outlined in black. Note that only the right or left side of each frontal section is shown. Images taken of the embryos' right side (I, K, L) have been mirrored to match images taken of the left side. Images A-F, H-L taken at 10X, G taken at 5X. Mx: maxilla, Md: mandible, T: tongue.

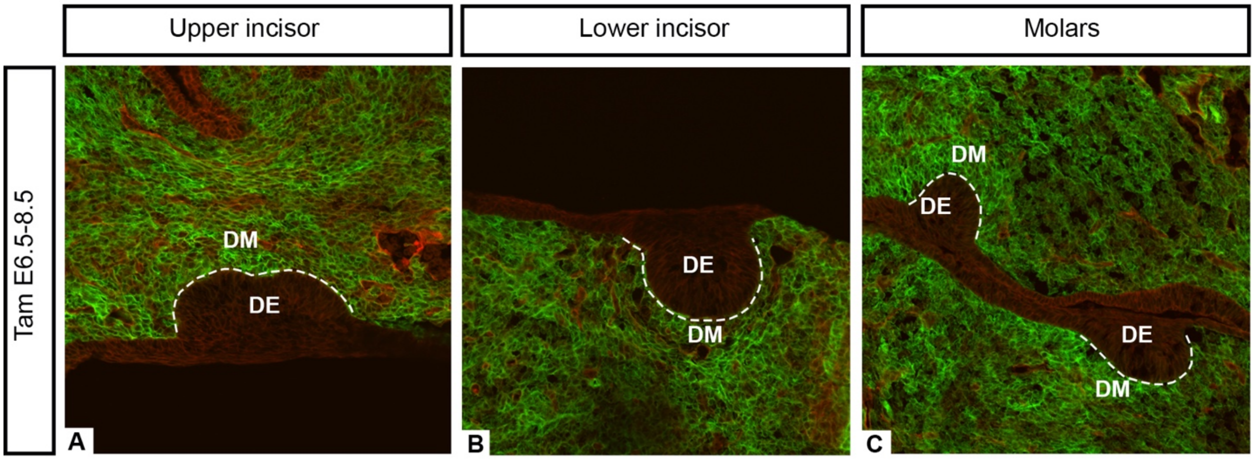


**Supplementary Figure 3. Frontal sections through bud stage (E13.5) incisors (A,B) and molars (C) showing CNCC-derived mesenchyme cells (EGFP-positive) labeled for three consecutive days via tamoxifen administration at E6.5-8.5 in *Sox10-iCre/ER*$^{T2}$*;R26R*$^{mTmG/+}$ or *Sox10-iCre/ER*$^{T2}$*;R26R*$^{mTmG/mTmG}$ embryos.** The majority of dental mesenchyme cells are labeled with EGFP indicating that between ~E6.5 and 8.5 NCCs contribute extensively to incisor and molar tooth germs. The dental epithelium is outlined in white. Images taken at 20X. DE: dental epithelium, DM: dental mesenchyme.

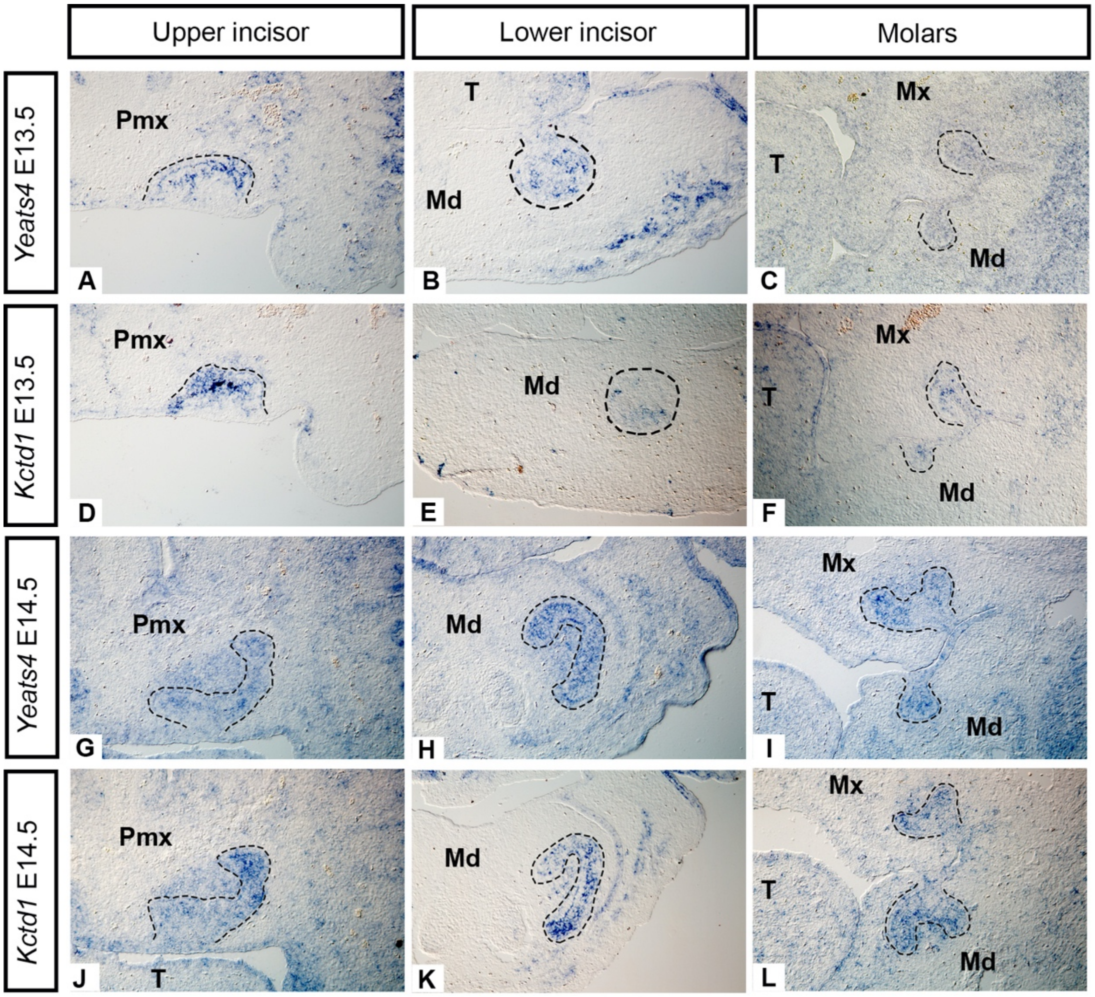


**Supplementary Figure 4. mRNA expression of *Yeats4,* and *Kctd1* at the late bud stage (E13.5, A-F) and cap stage (E14.5, G-L).** mRNA transcripts detected by *in-situ* hybridization on frontal cryosections from the upper incisor (left column), lower incisor (middle column), and molars (right column). Note that for both *Kctd1* and *Yeats4* a stronger signal was detected at E14.5 than E13.5, predominantly in the dental epithelium but expression is also present in the mesenchyme at the cap stage. At 14.5, the expression patterns of *Yeats4* and *Kctd1* are highly similar to that of *Tfap2a* (Figure 2 M-O). Note that only the right or left side of each frontal section is shown. Images taken of the embryos' right side (C, F, E, K) have been mirrored to match images taken of the left side. All images taken at 10X. Pmx: premaxilla, Mx: maxilla, Md: mandible, T: tongue.

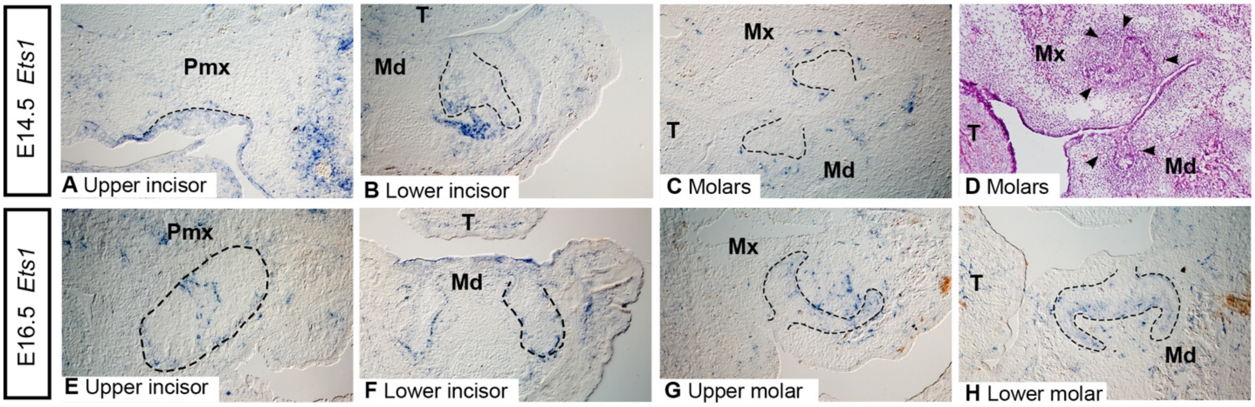


**Supplementary Figure 5. Sparse, punctate expression of *Ets1* at the cap (E14.5) and early bell (E16.5) stages revealed by *in situ* hybridization.** Cryosections in the frontal plane showing localization of mRNA transcripts of *Ets1* in incisor (A, B, E, F) and molar tooth germs (C, G, H). Note the similarity in the distribution of *Ets1* expressing cells (C) and red blood cells (D, arrowheads). Note that only the right or left side of each frontal section is shown. Images taken of the embryos' right side (E, G, H) have been mirrored to match images taken of the left side. Images taken at 10X. Pmx: premaxilla, Mx: maxilla, Md: mandible, T: tongue.

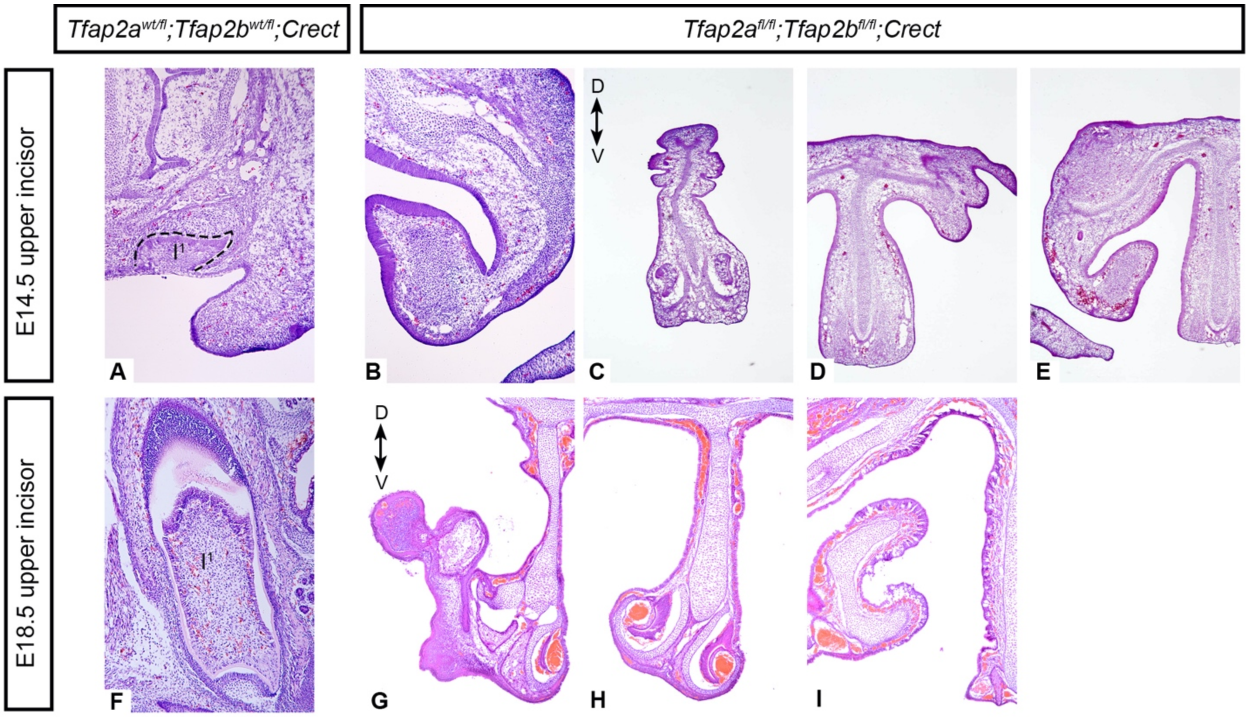


**Supplementary Figure 6. Upper incisors are absent in some *Tfap2a$^{fl/fl}$;Tfap2b$^{fl/fl}$;Crect* embryos at E14.5 (B-E) and E18.5 (G-I).** Histological sections in the frontal plane were stained with hematoxylin and eosin. Sections C-E and G-I are arranged with anterior-most sections on the left and more posterior sections on the right from the same individual embryos, respectively. Sections A (control) and B (mutant) are from comparable sectional planes along the anterior-posterior axis of the head. Note the right side of the E14.5 sections (A-E) and the left side of the E18.5 sections (F-I) are shown. Images A, B, F taken at 10X, C-E, G-I taken at 5X.

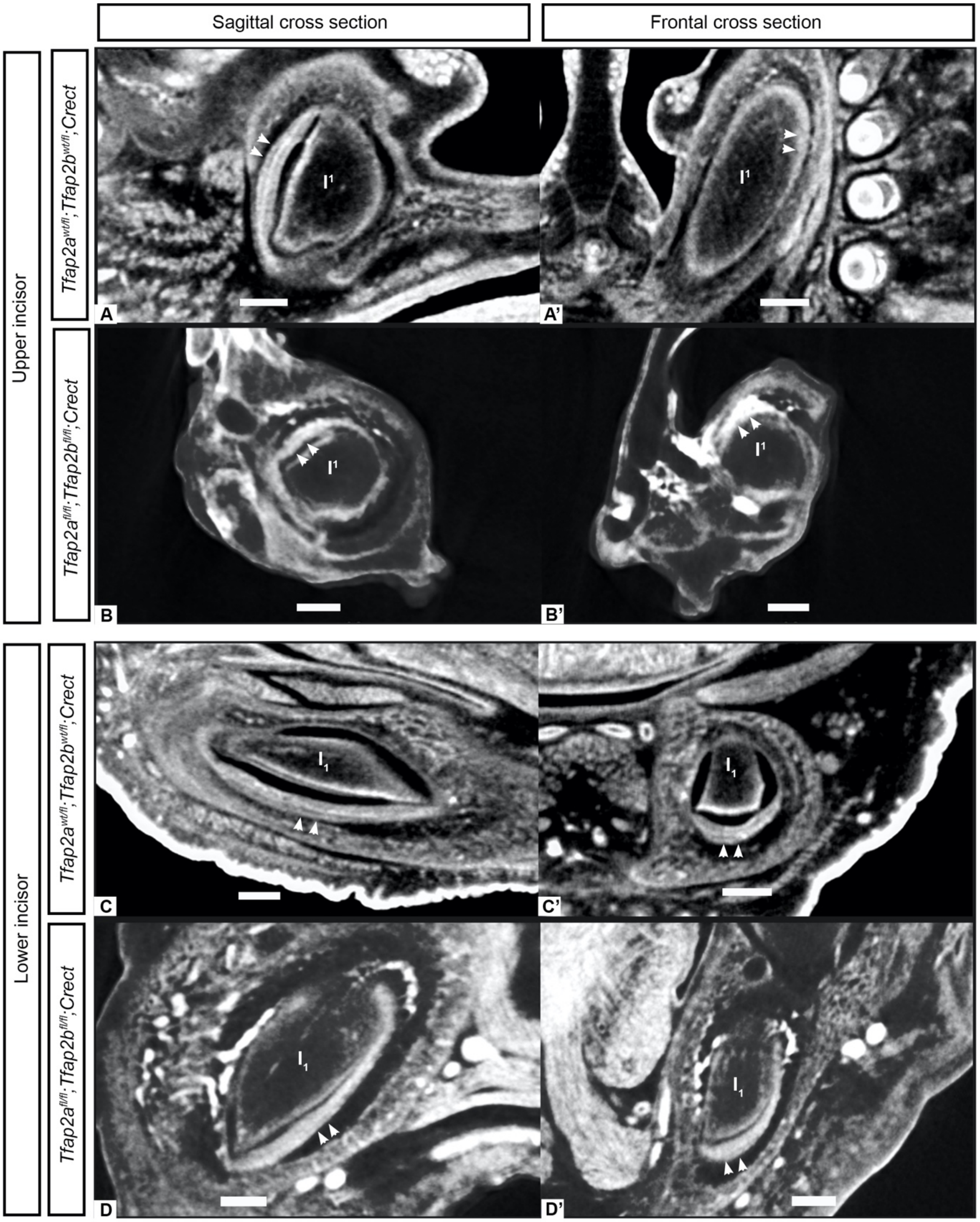


**Supplementary Figure 7. Comparison of upper (A-B) and lower (C-D) incisors in control (A, A', C, C') and *Tfap2a*$^{fl/fl}$*;Tfap2b*$^{fl/fl}$*;Crect* mutant (B, B', D, D') embryos at E18.5.** Sagittal (A-D) and lateral (A'-D') sections from µCT data show the left upper or lower incisors, respectively. Note the whiter regions along the periphery of the tooth are the most dense and correspond to enamel (white arrowheads). Scale bars are 0.2mm.

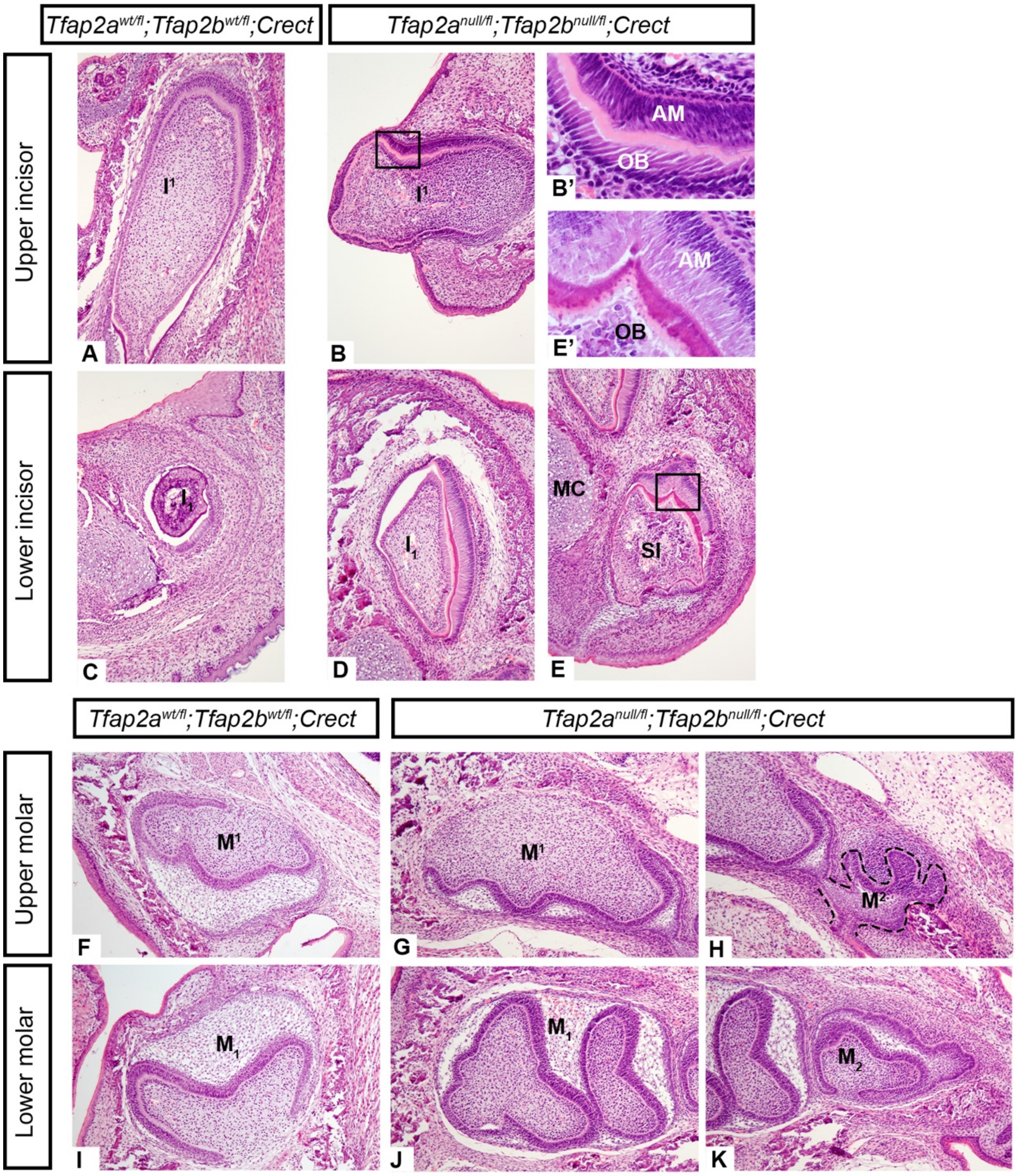


**Supplementary Figure 8. Comparison of the dentition in control (A, C, F, I) and mutant (*Tfap2a*$^{null/fl}$*;Tfap2b*$^{null/fl}$*;Crect*) embryos (B, D, E, G, H, J, K) show that upper incisors are present in the mutant (B) though the tooth is oriented horizontally as opposed to vertically in the control (A).** Hematoxylin and eosin stained cryosections in the frontal plane showing that both upper and lower incisors are present in the control (A, C) and in the mutant (B, D). A ventral supernumerary incisor (E) is also present in the lower dentition of mutant embryos. Ameloblasts and odontoblasts are also present in the upper incisor (B, B') and in the supernumerary lower incisor (E, E') of the mutant. Molars in the mutant (G, H, J, K) appear similar to those of the control (F, I). Subtle differences in the shape of the mutant molars (G, J, H, K) and the apparent medial displacement of $M^2/_2$ relative to $M^1/_1$ (H, K) compared with those of the control (F, I) are due to a slightly offset plane of section resulting from the cleft palate and the mandible which is both clefted and downturned in the mutants (data not shown), as seen in the *Tfap2a*$^{fl/fl;}$*Tfap2b*$^{fl/fl}$*;Crect* mutants (Figure 4). Note that only the right or left side of each frontal section is shown. Images taken of the embryos' right side (A, C) have been mirrored to match images taken of the left side. Images A-K taken at 10X, B', E' taken at 40X. SI: supernumerary incisor, MC: Meckel's cartilage, OB: odontoblasts, AM: ameloblasts.

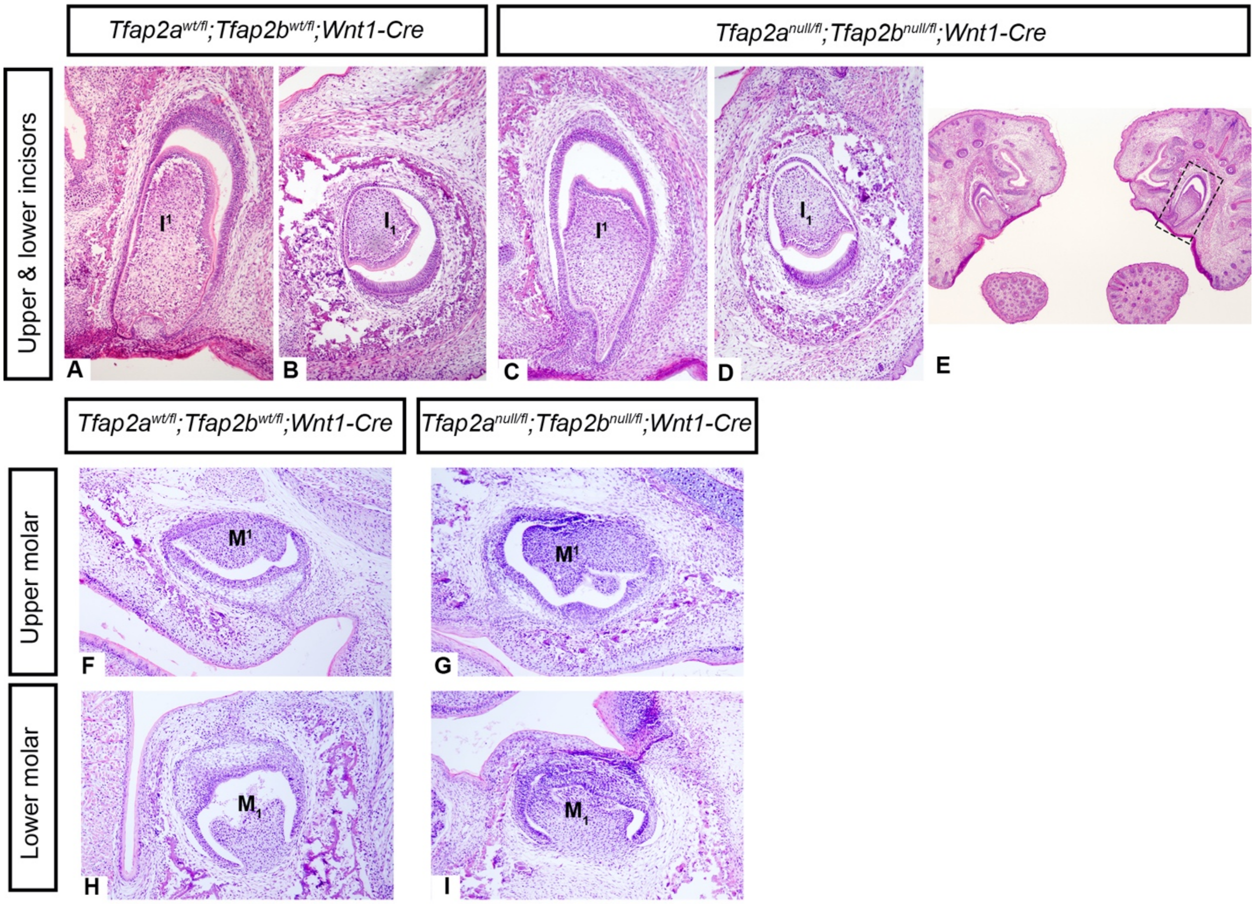


**Supplementary Figure 9. Teeth develop normally through the bell stage (E18.5) in mice lacking *Tfap2a* and *Tfap2b* in the neural crest derived mesenchyme (*$Tfap2a^{null/fl}$;$Tfap2b^{null/fl}$; Wnt1-Cre*), despite clefting in the palate and mandible (E).** Hematoxylin and eosin stained cryosections in the frontal plane showing that upper incisors (A, C) and lower incisors (B, D) appear similar in the mutant (C, D) compared to the control (A, B). Box in E surrounds upper incisor shown in C. Development of upper molars (F, G) and lower molars (H, I) also appears unperturbed in the mutant (G, I). Note that only the right or left side of each frontal section is shown. Images taken of the embryos' right side (A, B, F, H) have been mirrored to match images taken of the left side. Images A-D, F-I taken at 10X, image E taken at 5X.

**Supplementary Table 1. Primer sequences for genotyping mice and embryos.** ND: not detected, NA: not applicable. Primers for *Sox10-iCre/ER*$^{T2}$ mice obtained from: McKenzie, I.A., Ohayon, D., Li, H., de Faria, J.P., Emery, B., Tohyama, K., Richardson, W.D., 2014. Motor skill learning requires active central myelination. Science 346, 318–322. https://doi.org/10.1126/science.1254960

| Target | Primer Sequence 5'- 3' | Tm (°C) | Wild Type | Conditional | Mutant |
|---|---|---|---|---|---|
| *Sox10-iCre/ER*$^{T2}$ Fwd (McKenzie et al. 2014) | TTGCGATGGGAGAGTCTGAC | 64 | ND | NA | 742 bp |
| *Sox10-iCre/ER*$^{T2}$ Rev (McKenzie et al. 2014) | AGGTACAGGAGGTAGTCCCTC | | | | |
| *R26R*$^{mTmG/mTmG}$ Fwd (Endogenous) (JAX 8545) | AAAGTCGCTCTGAGTTGTTAT | 64 | 558 bp | NA | 337 bp |
| *R26R*$^{mTmG/mTmG}$ Rev 1 (Endogenous) (JAX 8546) | GGAGCGGGAGAAATGGATATG | | | | |
| *R26R*$^{mTmG/mTmG}$ Rev 2 (Mutant) (JAX 7320) | TCAATGGGCGGGGGTCGTT | | | | |
| *Tfap2a* | GCTCTCTCTTTTCCTGCCTTGGAACCATGACCCTCAG | 70 | 230 bp | 265 bp | ND |
| *Tfap2a* | GTAACGTTGGCAGCTTTACGTCTCCCTGCTGGC | | | | |
| *Tfap2b* | GTCTGTTTAGAACCTGGCTCAGCCAGAGGCTGG | 70 | 210 bp | 300 bp | ND |
| *Tfap2b* | CCCGAGCTAAGTGAACAGCTTCCCCTGTAAGGAGAGC | | | | |
| *Wnt1-Cre* | GGAGCGTCCACTGGAGTCCAGGTCCTCTGGTTG | 70 | ~310 bp | NA | ~220 bp |
| *Wnt1-Cre* | CAAACTAGGGTCATTAGACTTACAAGGCATGTG | | | | |
| *Wnt1-Cre* | CGCCCAATACCCTACTCTTCCGGAGGAAAATGTC | | | | |
| *Crect* | CCTCACTGATCCACATATGTCCTTCCGAAAGCTGC | 68 | 586 bp | NA | 385 bp |
| *Crect* | GATGCTAGAAAGCTGAGGCTGGGCTTAGCTTGCTAGGC | | | | |
| *Crect* | CTACGCCGCGAACTTGCTTCTAGAGCG | | | | |

**Supplementary Table 2. Dental phenotypes associated with different genotypes from *Crect* and *Wnt1-Cre* crosses.**

| Mutant or Control | Genotype | Stage | Affected Tissue | N | Dental Phenotype |
|---|---|---|---|---|---|
| Mutant | *Tfap2a*$^{fl/fl}$;*Tfap2b*$^{fl/fl}$; *Crect* | 18.5 (bell) | Ectoderm (including dental epithelium) | 4 | Duplicated lower incisors (N=2/4)<br>Ventrally-curved lower incisors (N=2/4)<br>Upper incisors absent (3/4)<br>Molars mesiodistally shortened (N=1) |
| Mutant | *Tfap2a*$^{fl/fl}$;*Tfap2b*$^{fl/fl}$;*Crect* | 14.5 (cap) | Ectoderm (including dental epithelium) | 3 | Duplicated lower incisors (N=3/3)<br>Upper incisors absent (N=3/3) |
| Mutant | *Tfap2a*$^{fl/null}$;*Tfap2b*$^{fl/null}$;*Crect* | 18.5 (bell) | Ectoderm (including dental epithelium); *Cre*-negative tissues heterozygous for *Tfap2a/Tfap2b* | 2 | Duplicated lower incisors (N=2/2)<br>One upper incisor absent (N=2/2) |
| Control | *Tfap2a*$^{fl/wt}$;*Tfap2b*$^{fl/wt}$;*Crect* | 18.5 (bell) | None | 3 | Wild-type incisors and molars |
| Control | *Tfap2a*$^{fl/wt}$;*Tfap2b*$^{fl/wt}$;*Crect* | 14.5 (cap) | None | 3 | Wild-type incisors and molars |
| Mutant | *Tfap2a*$^{fl/fl}$;*Tfap2b*$^{fl/fl}$;*Wnt1-Cre* | 18.5 (bell) | Neural crest-derived mesenchyme | 3 | Wild-type incisors (N=3) and molars (N=2). Molars mesiodistally shortened (N=1). Cleft in midface (N=3) |
| Mutant | *Tfap2a*$^{fl/fl}$;*Tfap2b*$^{fl/fl}$;*Wnt1-Cre* | 14.5 (cap) | Neural crest-derived mesenchyme | 3 | Wild-type incisors and molars |
| Mutant | *Tfap2a*$^{fl/null}$;*Tfap2b*$^{fl/null}$;*Wnt1-Cre* | 17.5-18.5 (bell) | Neural crest-derived mesenchyme; *Cre*-negative tissues heterozygous for *Tfap2a/Tfap2b* | 3 | Wild-type incisors (N=3) and molars (N=2). Unknown if molars are mesiodistally shortened.<br>Cleft in midface and mandible (N=3/3) |
| Control | *Tfap2a*$^{fl/wt}$;*Tfap2b*$^{fl/wt}$;*Wnt1-Cre* | 17.5-18.5 (bell) | None | 3 | Wild-type incisors and molars |
| Control | *Tfap2a*$^{fl/wt}$;*Tfap2b*$^{fl/wt}$;*Wnt1-Cre* | 14.5 (cap) | None | 3 | Wild-type incisors and molars |

**Supplementary Table 3. Primers used to amplify DNA sequences of target genes used for making mRNA probes for *in situ* hybridization.**

| Primer Name | Primer Sequence (5'-3') | Tm (°C) | Amplicon (bp) |
|---|---|---|---|
| *Tfap2a* Forward | GACCGTCACGACGGCACCAG | 64 | 433 |
| *Tfap2a* Reverse | GGACGTCCTCGATGGCGTGAG | | |
| *Tfap2b* Forward | GCCTTGCTCTTACTGTGCAG | 65 | 454 |
| *Tfap2b* Reverse | CTATCTAGCTGCCCCTTCGC | | |
| *Ets1* Forward | GCTACGGTATCGAGCATGCTC | 64 | 431 |
| *Ets1* Reverse | CCAGGCACATGTTGTCTGGAG | | |
| *Kctd1* Forward | GCATGTACTTCTGCACGCGAG | 61 | 282 |
| *Kctd1* Reverse | GCCGCTGGAAAAACGCCTTA | | |
| *Yeats4* Forward | CAGAACTTGAAGTGAAAACCAG | 54 | 315 |
| *Yeats4* Reverse | TGGAGTCCTCTCTGAGAAAG | | |

**Supplementary Table 4. Upper and lower first molar measurements of wild mice and μCT-scanned embryos.** Measurements (in mm) of molar occlusal surfaces of wild mice, *Mus musculus,* were obtained from Csanady, A., Mosansky, L., 2018. Skull morphometry and sexual size dimorphism in Mus musculus from Slovakia. North-Western Journal of Zoology 14, 102–106. Measurements of control and mutant molars were obtained from μCT scanned embryos from this study. Standard deviations for the mean measurements from the wild mice are also listed. The largest standard deviation (0.08, indicated with an asterisk) was used to estimate minimum and maximum values for molar length and width for the embryos examined in this study.

| **Tooth measurement** | ***Mus musculus*** | ***Tfap2a$^{fl/wt}$*; *Tfap2b$^{fl/wt}$*; *Crect* (Control)** | ***Tfap2a$^{fl/fl}$*; *Tfap2b$^{fl/fl}$*; *Crect* (Mutant)** | ***Tfap2a$^{fl/wt}$*; *Tfap2b$^{fl/wt}$*; *Wnt1Cre* (Control)** | ***Tfap2a$^{fl/fl}$*; *Tfap2b$^{fl/fl}$*; *Wnt1Cre* (Mutant)** |
|---|---|---|---|---|---|
| Lower M1 mean length (mm) | 1.34 +/- 0.06 | 1.043 | 0.857 | 1.157 | 0.718 |
| Lower M1 mean width (mm) | 0.78 +/- 0.03 | 0.577 | 0.558 | 0.590 | 0.508 |
| Upper M1 mean length (mm) | 1.61 +/- 0.08 * | 1.210 | 0.953 | 1.297 | 0.926 |
| Upper M1 mean width (mm) | 0.98 +/- 0.04 | 0.636 | 0.613 | 0.665 | 0.611 |
